# EIPR1 controls dense-core vesicle cargo retention and EARP complex localization in insulin-secreting cells

**DOI:** 10.1101/374488

**Authors:** Irini Topalidou, Jérôme Cattin-Ortolá, Blake Hummer, Cedric S. Asensio, Michael Ailion

## Abstract

Dense-core vesicles (DCVs) are secretory vesicles found in neurons and endocrine cells. DCVs package and release cargos including neuropeptides, biogenic amines, and peptide hormones. We recently identified the endosome-associated recycling protein (EARP) complex and the EARP-interacting protein EIPR-1 as proteins important for controlling levels of DCV cargos in *C. elegans* neurons. Here we determine the role of mammalian EIPR1 in insulinoma cells. We find that in *Eipr1* KO cells, there is reduced insulin secretion, and mature DCV cargos such as insulin and carboxypeptidase E (CPE) accumulate near the trans-Golgi network and are not retained in mature DCVs in the cell periphery. In addition, we find that EIPR1 is required for the stability of the EARP complex subunits and for the localization of EARP and its association with membranes, but EIPR1 does not affect localization or function of the related Golgi-associated retrograde protein (GARP) complex. EARP is localized to two distinct compartments related to its function: an endosomal compartment and a DCV biogenesis-related compartment. We propose that EIPR1 functions with EARP to control both endocytic recycling and DCV maturation.

## Introduction

Dense-core vesicles (DCVs) are regulated secretory vesicles found in neurons and endocrine cells, where they are also called secretory granules. DCVs package several types of cargo, including neuropeptides and peptide hormones, for release at the cell membrane (Gondré-Lewis *et al*., 2012). The secreted cargos modulate a variety of processes including development, growth, glucose metabolism, and mental state. DCVs are generated at the trans-Golgi network (TGN) in a process that includes correct sorting of cargo and acquisition of proper compartmental identity. Because DCVs are not regenerated locally at release sites, the DCV pool needs to be continuously supplied by the TGN.

Genetic studies in the nematode *C. elegans* have identified several molecules that function in DCV biogenesis, including the endosome-associated recycling protein (EARP) complex and the EARP-interacting protein EIPR-1, a WD40 domain protein that interacts with EARP (Edwards *et al*., 2009; Sumakovic *et al*., 2009; Mesa *et al*., 2011; Yu, Wang, Jiu, *et al*., 2011; Hannemann *et al*., 2012; Ailion *et al*., 2014; Topalidou *et al*., 2016). The EARP complex is structurally similar to the Golgi-associated retrograde protein (GARP) complex. EARP shares the VPS51, VPS52, and VPS53 subunits with the GARP complex, but uses VPS50 instead of VPS54 as the fourth subunit (Gillingham *et al*., 2014; Schindler, Chen, *et al*., 2015). Whereas GARP functions in retrograde trafficking from endosomes to the TGN (Conibear and Stevens, 2000; Conibear *et al*., 2003; Pérez-Victoria *et al*., 2008; Pérez-Victoria and Bonifacino, 2009; Perez-Victoria *et al*., 2010), EARP was shown to act in recycling cargos from endosomes back to the plasma membrane (Schindler, Chen, *et al*., 2015). In *C. elegans*, the EARP complex and EIPR-1 were shown to be required for controlling levels of DCV cargos (Topalidou *et al*., 2016). The VPS50 subunit of the EARP complex was also shown to be required for the maturation of DCV cargos and DCV acidification (Paquin *et al*., 2016).

EIPR1 physically interacts with the EARP complex in rat insulinoma cells (Topalidou *et al*., 2016). EIPR1 (also named TSSC1) was also independently identified as a physical interactor and functional partner of both the GARP and EARP complexes in human cell lines (Gershlick *et al*., 2016). Moreover, two mass spectrometry interactome data sets identified EIPR1 as an interactor of EARP subunits in human HEK293T and HeLa cells (Hein *et al*., 2015; Huttlin *et al*., 2015). WD40 domain proteins like EIPR1 often act as scaffolds for the assembly of protein complexes (Stirnimann, Petsalaki, *et al*., 2010). Though EIPR1 interacts with EARP, it has not been determined whether EIPR1 is required for the localization or stability of the EARP complex. Fluorescence recovery after photobleaching in *Eipr1* knockdown cells showed that EIPR1 is required for efficient recruitment of GARP to the TGN (Gershlick *et al*., 2016).

Here we investigated the role of mammalian EIPR1 in DCV function and EARP complex formation using insulin-secreting insulinoma cells. Specifically, we used *Eipr1* knockout and rescue experiments to demonstrate that EIPR1 controls proper insulin distribution and secretion, and retention of cargo in mature DCVs. We also found that EIPR1 is required for the stability of the EARP complex subunits and for proper localization and association of EARP with membranes. Finally, we found that EARP localizes to two distinct compartments relevant to its functions in endocytic recycling and DCV maturation.

## Results

### EIPR1 is required for insulin secretion

The *C. elegans* WD40 domain protein EIPR-1 is needed for dense-core vesicle (DCV) cargo trafficking in *C. elegans* neurons (Topalidou *et al*., 2016). To investigate the role of EIPR1 in the trafficking of mammalian DCV cargo in endocrine cells, we generated *Eipr1* knockout (KO) insulinoma 832/13 cells using the CRISPR technology by inserting a puromycin cassette in the first exon of *Eipr1* (Figure 1A and S1A,B). We identified positive clones by PCR (Figure S1C). To confirm that EIPR1 is lost in the *Eipr1* KO line, we analyzed the cells for EIPR1 expression by Western blot. Wild-type (WT) cells displayed a band at around 45 kD, the approximate molecular weight of EIPR1, which was missing from *Eipr1* KO cells (Figure 1B).

**Figure 1.**
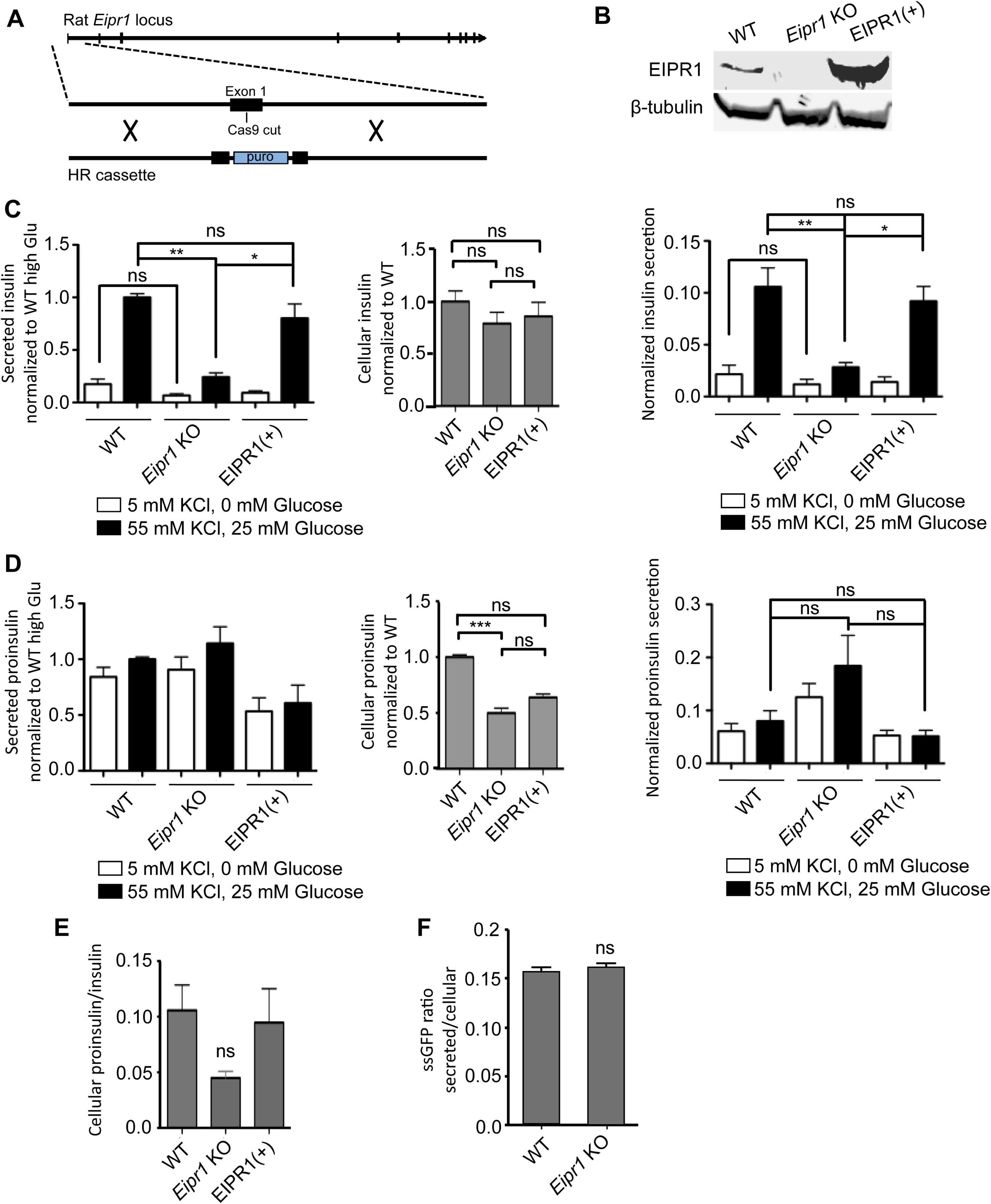
Insulin secretion is reduced in *Eipr1* KO cells. (A) Strategy used to create the *Eipr1* KO 832/13 cell line. Cas9 was targeted to cut in the first exon of the rat *Eipr1* locus and homologous recombination (HR) was used to insert a puromycin cassette. (B) *Eipr1* KO cells do not express wild type EIPR1. Protein extracts from 832/13 (WT), *Eipr1* KO 832/13 (Eipr1KO), and *Eipr1* KO 832/13 cells expressing a wild type *Eipr1* cDNA (EIPR1(+)) were blotted with an EIPR1 antibody. β-tubulin served as a loading control. (C) (Left panel) Insulin secretion under resting (5 mM KCl, 0 mM glucose) and stimulating conditions (55 mM KCl, 25 mM glucose) from 832/13 cells (WT), *Eipr1* KO 832/13 cells (Eipr1KO), and an *Eipr1* KO stable line expressing wild type *Eipr1* (EIPR1(+)). All values were normalized to the value of the WT under stimulating conditions. n= 7, * p<0.05, ** p<0.01, ns p>0.05, error bars = SEM. (Middle panel) Total insulin content in WT, *Eipr1* KO, and EIPR1(+) cells. All values were normalized to the WT. n= 5-7; ns p>0.05, error bars = SEM. (Right panel) Insulin secretion normalized to insulin content under resting (5 mM KCl, 0 mM glucose) and stimulating conditions (55 mM KCl, 25 mM glucose) from WT, *Eipr1* KO, and EIPR1(+) cells. n= 5-7, * p<0.05, ** p<0.01, ns p>0.05, error bars = SEM. We performed three biological replicates. For each replicate, the same cells were used to determine the amount of insulin secreted under resting conditions, stimulating conditions, and the amount of total cellular insulin. (D) (Left panel) Proinsulin secretion under resting (5 mM KCl, 0 mM glucose) and stimulating conditions (55 mM KCl, 25 mM glucose) from WT, *Eipr1* KO, and EIPR1(+) cells. All values were normalized to the value of the WT under stimulating conditions. n= 6, error bars = SEM. (Middle panel) Total proinsulin content in WT, *Eipr1* KO, and EIPR1(+) cells. All values were normalized to the WT. n= 6, *** p<0.001, ns p>0.05, error bars = SEM. (Right panel) Proinsulin secretion normalized to proinsulin content under resting (5 mM KCl, 0 mM glucose) and stimulating conditions (55 mM KCl, 25 mM glucose) from WT, *Eipr1* KO, and EIPR1(+) cells. All values were normalized to the WT under stimulating conditions. n= 6, ns p>0.05, error bars = SEM. We performed three biological replicates. (E) Ratio of total cellular proinsulin to total insulin. n= 4-6, ns p>0.05, error bars = SEM. (F) The absence of EIPR1 does not affect the constitutive secretory pathway. *Eipr1* KO 832/12 cells secrete normal levels of ssGFP (GFP fused to a signal peptide at its N-terminus). Values of secreted GFP were normalized to total. n=6, error bars = SEM. The data shown were combined from two independent experiments with similar results. The data shown for the WT are the same shown in Figure 1D of (Cattin-Ortolá, Topalidou *et al*., 2019) since these experiments were run in parallel with the same WT control.

To examine whether EIPR1 is needed for DCV cargo trafficking in insulinoma cells, we measured insulin secretion of WT and *Eipr1* KO cells under resting (5 mM KCl, 0 mM glucose) and stimulating (55 mM KCl, 25 mM glucose) conditions. Insulin secretion under stimulating conditions was lower in *Eipr1* KO cells than in WT cells (stimulated secretion was reduced to ∼ 25% of WT, Figure 1C, left panel). Insulin secretion under resting conditions was also lower, but the difference was not statistically significant (Figure 1C, left panel). To verify that the effects were due to loss of EIPR1, we introduced a wild type *Eipr1* cDNA into the *Eipr1* KO cells (Figure 1B). Expression of wild type EIPR1 in *Eipr1* KO cells rescued the stimulated insulin secretion defect of the *Eipr1* KO line, confirming that this defect is due to loss of EIPR1 (Figure 1C, left panel, EIPR1(+)).

Defective insulin secretion can be due to reduced insulin content. Total insulin content of *Eipr1* KO was slightly reduced (to ∼ 80% of WT, Figure 1C, middle panel), but the difference was not statistically significant. After normalizing insulin secretion to total insulin content, we found that insulin secretion under stimulating conditions was still significantly reduced in the *Eipr1* KO line (∼30% of WT, Figure 1C, right panel).

The observed decrease in insulin secretion in the *Eipr1* KO could be due to a defect in the processing of proinsulin to insulin. Thus, we measured the total and secreted levels of proinsulin. Although secretion of proinsulin was not altered in *Eipr1* KO (Figure 1D, left panel), total proinsulin content was reduced (∼50% of WT, Figure 1D, middle panel). To examine whether the reduced level of total proinsulin is due to a transcription defect, we performed quantitative RT-PCR. No difference was observed in the level of proinsulin mRNA in *Eipr1* KO cells (Figure S2A). The reduction in total proinsulin was not significantly rescued in EIPR1(+) cells (Figure 1D, middle), so it may be due to an EIPR1-independent defect of the *Eipr1* KO cell line. The ratio of total cellular proinsulin/insulin was also reduced in the *Eipr1* KO, but the effect was not statistically significant (Figure 1E). These results suggest that proinsulin processing and secretion are not strongly affected by the absence of EIPR1.

One possible explanation for the reduced insulin secretion in *Eipr1* KO cells is that exocytosis is impaired. We counted exocytotic events in WT and *Eipr1* KO cells expressing the neuropeptide NPY tagged to pH-sensitive GFP (NPY::pHluorin). Exocytotic events were reduced in *Eipr1* KO cells under stimulating conditions (Figure S3), suggesting that *Eipr1* KO cells have an exocytosis defect. However, a caveat to this experiment is that this assay may have underestimated the true exocytosis rate if it was not possible to detect DCVs carrying reduced amounts of NPY::pHluorin. As *Eipr1* KO cells appear to carry reduced amounts of cargos in mature DCVs (see below), it is possible that DCV exocytotic events in this mutant line were more difficult to detect.

To investigate whether EIPR1 is needed for constitutive secretion we measured the secretion of GFP fused to a signal peptide (ssGFP) under resting conditions (Hummer *et al*., 2017). We found that secretion of ssGFP in *Eipr1* KO cells is similar to WT, suggesting that EIPR1 is not needed for constitutive secretion (Figure 1F).

### EIPR1 is required for the normal cellular distribution of insulin

To further investigate the role of EIPR1 in DCV cargo trafficking, we examined the subcellular localization of insulin by immunostaining. In wild type cells, insulin is detected as puncta spread throughout the cytoplasm (Figure 2A). By contrast, in *Eipr1* KO cells, insulin is localized primarily to a perinuclear region that partially overlaps with the *trans*-Golgi marker TGN38 (Figure 2A,B). This phenotype was rescued in *Eipr1* KO cells that stably express wild type EIPR1 (Figure 2A,B). The ratio of TGN38-localized insulin to the level of insulin in the cytoplasm was increased in *Eipr1* KO cells (Figure 2C), suggesting accumulation of insulin near the TGN, decreased insulin in the periphery of the cell, or both. To distinguish between these possibilities, we quantified the levels of insulin localized near the TGN and in the cytoplasm. We found that *Eipr1* KO cells have a small increase in insulin near the TGN (Figure 2D) and a decrease in insulin in the cell periphery (Figure 2E). The reduction in insulin in the cell periphery might contribute to the reduction in insulin secretion in *Eipr1* KO cells under stimulating conditions and reduced frequency of detectable exocytotic events (Figure 1C and S3).

**Figure 2.**
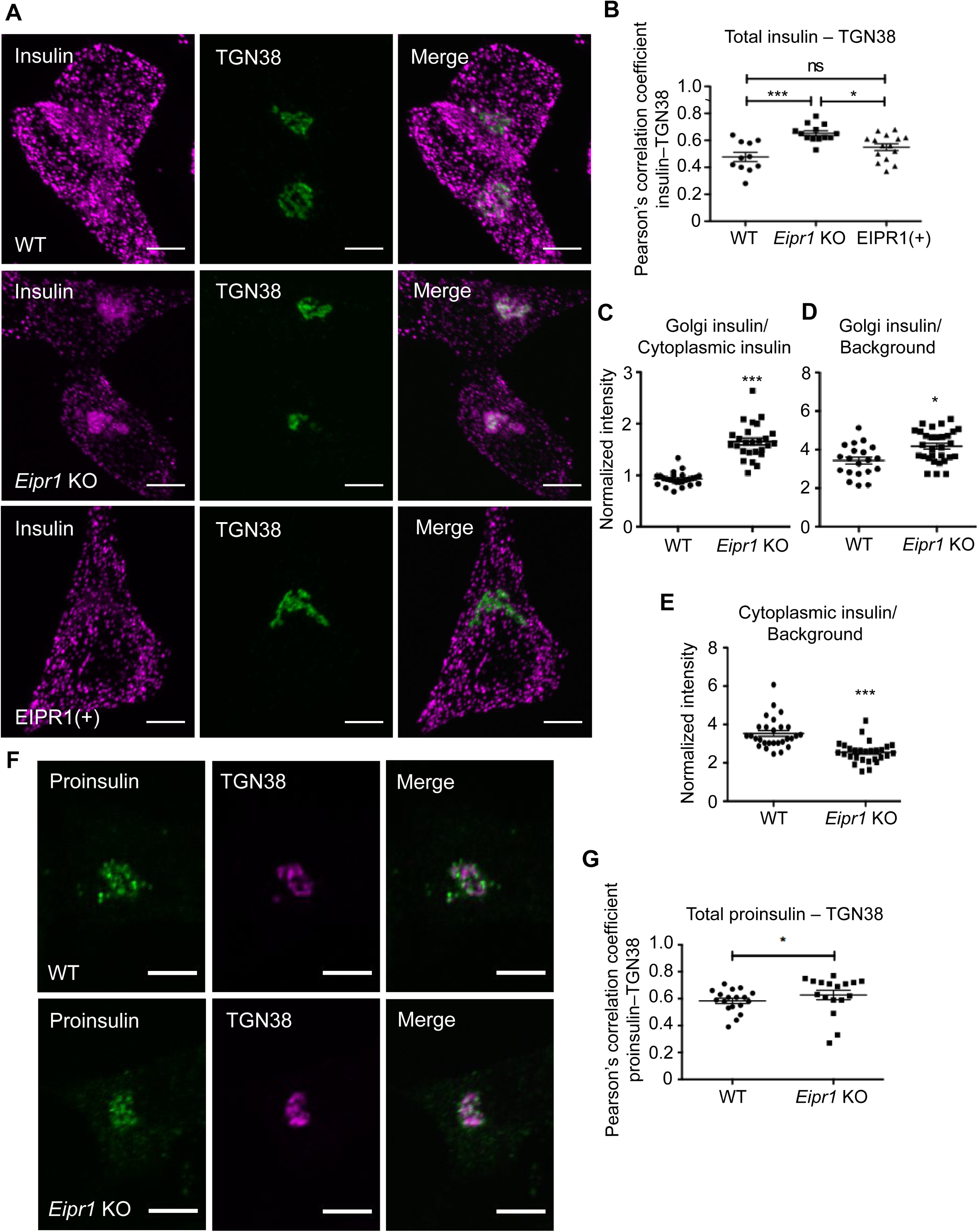
Insulin localization is disrupted in *Eipr1* KO cells. (A) Representative images of 832/13 (WT), *Eipr1* KO 832/13 (Eipr1KO), and *Eipr1* KO 832/13 cells expressing wild type *Eipr1* (EIPR1(+)) costained for endogenous insulin and TGN38. In WT and EIPR1(+) cells, insulin is spread throughout the cytoplasm, but in *Eipr1* KO cells insulin accumulates in a perinuclear region that partially overlaps with TGN38. The experiment was repeated three times and the experimenter was blinded to the genotypes of the stained cells. Scale bars: 5 μm. (B) Pearson’s correlation coefficient was measured to quantify the localization between insulin and the TGN marker TGN38. n=11 for WT, n=13 for *Eipr1* KO and n=15 for EIPR1(+), *** p<0.001, * p<0.05, ns p>0.05, error bars = SEM. The experiment was repeated three times. (C) *Eipr1* KO cells have an increased Golgi/cytoplasmic ratio of insulin relative to wild type cells. Fluorescence of a region of interest that includes the TGN divided by the fluorescence of a region of the same size in the cytoplasm, in WT and *Eipr1* KO. (n=25 for WT and *Eipr1* KO, error bars = SEM, ***p<0.001). The data shown for the WT are the same shown in Figure 2C of (Cattin-Ortolá, Topalidou *et al*., 2019) since these experiments were run in parallel with the same WT control. (D) *Eipr1* KO cells have a slightly increased amount of insulin localized at or near the TGN. Fluorescence of a region of interest that includes the TGN divided by the fluorescence of a region of the same size in the background, in WT and *Eipr1* KO. (n=20 for WT and n = 30 for *Eipr1* KO, error bars = SEM, *p<0.05). (E) *Eipr1* KO cells have decreased amount of insulin localized to the cytoplasm. Fluorescence of a region of interest in the cytoplasm divided by the fluorescence of a region of the same size in the background (n=28 for WT and *Eipr1* KO, error bars = SEM, ***p<0.001). (F) Representative images of 832/13 (WT) and *Eipr1* KO 832/13 (Eipr1KO) cells costained for endogenous proinsulin and TGN38. In both WT and *Eipr1* KO cells, proinsulin is localized in a perinuclear region that partially overlaps with TGN38. Scale bars: 5 μm. (G) Pearson’s correlation coefficient was measured to quantify the localization between proinsulin and the TGN marker TGN38. n=18 for WT and n=17 for *Eipr1* KO * p<0.05, error bars = SEM. The experiment was repeated three times.

We also examined the localization of proinsulin, a cargo of immature DCVs. Proinsulin is localized in a similar perinuclear region in both wild type and *Eipr1* KO cells, with perhaps a very slight shift towards tighter colocalization with TGN38 in the *Eipr1* KO (Figure 2F,G). Because insulin seems to be shifted to the TGN in *Eipr1* KO cells (Figure 2A-D), we wondered whether there might be increased acidification of the Golgi in these mutant cells which would allow processing of proinsulin into insulin in this compartment. To assay the pH of the Golgi, we measured the fluorescence of the pH-sensitive GFP variant pHluorin fused to the Golgi-localization domain of sialyltransferase (St6Gal1), a late-Golgi enzyme (Wong *et al*., 1992; Hummer *et al*., 2017). The pH of the Golgi is similar in *Eipr1* KO and WT cells (Figure S4), suggesting that the accumulation of insulin near the TGN in *Eipr1* KO cells is not due to proinsulin processing in the Golgi.

### EIPR1 is needed for the normal levels and distribution of mature DCV cargo

We next examined the levels of other known DCV cargos, such as the proprotein convertase 1/3 (PC1/3), proprotein convertase 2 (PC2), and carboxypetidase E (CPE) that mediate the processing of proinsulin to insulin (Smeekens *et al*., 1992; Naggert *et al*., 1995). These enzymes are themselves processed from their inactive pro forms to their active mature forms (Vindrola and Lindberg, 1992; Zhou and Lindberg, 1993; Song and Fricker, 1995; Muller and Lindberg, 1999).

The *Eipr1* KO cells had reduced levels of the processed form of PC1/3 but normal levels of PC2 (Figure 3A). Expression of wild type EIPR1 in *Eipr1* KO cells rescued the PC1/3 defect (Figure 3A). To examine whether the reduced levels of total PC1/3 are due to a transcription defect, we performed quantitative RT-PCR but found no difference between WT and *Eipr1* KO cells (Figure S2A).

**Figure 3.**
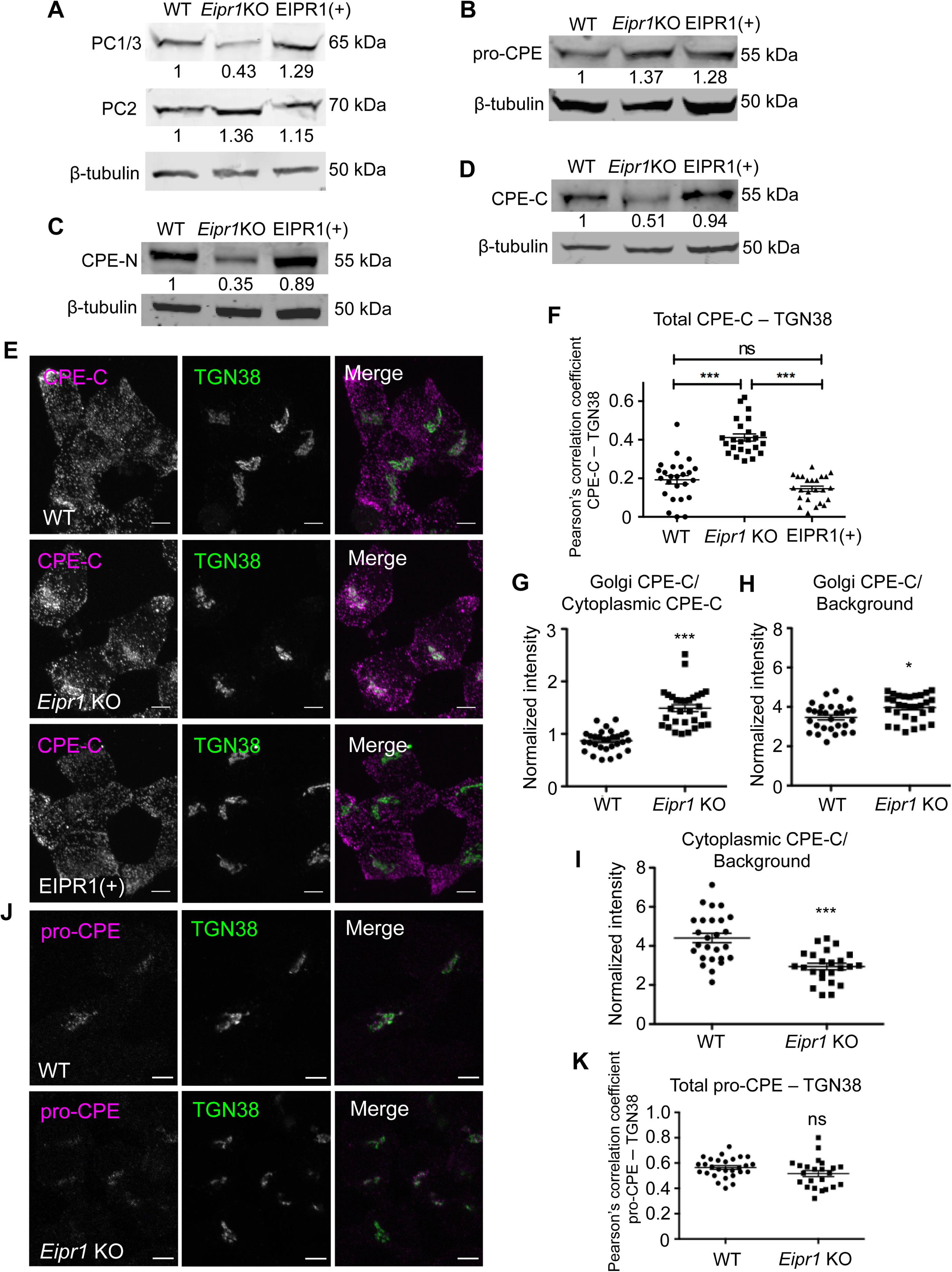
EIPR1 is required for normal levels and distribution of mature DCV cargos. (A) *Eipr1* KO cells have reduced levels of the processed form of the proprotein convertase PC1/3 but normal levels of PC2. Lysates from WT, *Eipr1* KO, and EIPR1(+) 832/13 cells were blotted with an antibody to PC1/3 or PC2. β-tubulin was used as a loading control. Shown are representative blots of three independent experiments with similar results. (B) The levels of the unprocessed form of CPE (pro-CPE) are not affected by the loss of EIPR1. Lysates from the indicated cell lines were blotted with an antibody to pro-CPE. The experiment was repeated three times with similar results. The data shown for the WT are the same shown in Figure 3B of (Cattin-Ortolá, Topalidou *et al*., 2019) since these experiments were run in parallel with the same WT control. (C), (D) *Eipr1* KO cells have reduced levels of the processed form of CPE. Detergent lysates from WT, *Eipr1* KO, and EIPR1(+) 832/13 cells were blotted with an antibody to the N-terminus of CPE (CPE-N) or the C-terminus of CPE (CPE-C). β-tubulin was used as a loading control. The experiment was repeated three times. The data shown for the WT in Figure 3C are the same shown in Figure 3C of (Cattin-Ortolá, Topalidou *et al*., 2019) since these experiments were run in parallel with the same WT control. (E) The mature processed form of CPE is localized at or near the TGN in *Eipr1* KO cells. Representative confocal images of WT, *Eipr1* KO, and EIPR1(+) cells costained with the CPE C-terminal antibody (CPE-C) and TGN38. Maximum intensity projections. Scale bars: 5 μm. The experiment was repeated three times with similar results. (F) Quantification of the colocalization between the mature form of CPE (CPE-C) and the TGN marker TGN38. Maximum intensity projection images were obtained and Pearson’s correlation coefficients were determined by drawing a line around each cell. (n=25 for WT; n=24 for *Eipr1* KO; n=24 for EIPR1(+), error bars = SEM, ***p<0.001, ns p>0.05). (G) *Eipr1* KO cells have an increased Golgi/cytoplasmic ratio of CPE relative to wild type cells. Fluorescence of a region of interest that includes the TGN divided by the fluorescence of a region of the same size in the cytoplasm, in WT and *Eipr1* KO. (n=30 for WT and *Eipr1* KO, error bars = SEM, ***p<0.001). The data shown for the WT are the same shown in Figure 3G of (Cattin-Ortolá, Topalidou *et al*., 2019) since these experiments were run in parallel with the same WT control. (H) *Eipr1* KO cells have an increased amount of mature CPE localized at the Golgi. Fluorescence of a region of interest at the Golgi divided by the fluorescence of a region of the same size in the background, in WT and *Eipr1* KO. (n=29 for WT and n = 31 for *Eipr1* KO, error bars = SEM, *p<0.05). (I) *Eipr1* KO cells have a decreased amount of insulin localized to the cytoplasm. Fluorescence of a region of interest in the cytoplasm divided by the fluorescence of a region of the same size in the background (n=26 for WT and n = 24 for *Eipr1* KO, error bars = SEM, ***p<0.001). (J) The unprocessed form of CPE (pro-CPE) is localized at or near the TGN in both WT and *Eipr1* KO cells. Representative confocal images of WT and *Eipr1* KO cells costained for pro-CPE antibody and TGN38. Single slices. Scale bars: 5 μm. The experiment was repeated twice. (K) Quantification of the colocalization between the unprocessed form of CPE (pro-CPE) and the TGN marker TGN38. Maximum intensity projection images were obtained and Pearson’s correlation coefficients were determined by drawing a line around each cell. (n=28 for WT; n=23 for *Eipr1* KO, error bars = SEM, ns p>0.05).

We measured the levels of CPE with antibodies that recognize the unprocessed CPE form that resides in the Golgi and immature DCVs (pro-CPE antibody), or the processed forms localized to mature DCVs (CPE-N and CPE-C antibodies) (Fricker *et al*., 1990, 1996). The *Eipr1* KO had normal levels of the unprocessed form of CPE (Figure 3B), but reduced levels of the processed form (Figure 3C,D). Expression of wild type EIPR1 in *Eipr1* KO cells rescued the mature CPE defects (Figure 3C,D). These results suggest that EIPR1 acts in a post-processing step and is required for the normal levels of mature DCV cargo.

We next examined the subcellular localization of CPE by immunostaining. In wild type cells, mature CPE was detected as puncta spread throughout the cytoplasm that almost fully colocalize with insulin (Figure 3E,F and S5). By contrast, in *Eipr1* KO cells, mature CPE was reduced in the cell periphery and increased in the TGN region (Figure 3E,F). This phenotype was rescued in *Eipr1* KO cells that stably expressed wild type EIPR1 (Figure 3E,F). As with insulin, the ratio of TGN38-localized mature CPE to mature CPE in the cytoplasm was increased in *Eipr1* KO cells (Figure 3G). *Eipr1* KO cells have slightly more mature CPE near the TGN (Figure 3H) and significantly less mature CPE in the cell periphery (Figure 3I). This loss of mature CPE from the cell periphery supports our finding that there are reduced levels of mature CPE in the *Eipr1* KO (Figure 3C,D). Pro-CPE was localized in a perinuclear region in both wild type and *Eipr1* KO cells (Figure 3J,K), similar to proinsulin (Figure 2F,G), consistent with EIPR1 acting in a post-processing step.

### Mature cargo is able to exit the TGN in *Eipr1* KO cells

To investigate whether the increased insulin and mature CPE at the TGN in *Eipr1* KO cells is due to mature DCV cargo being stuck at the TGN, we used a pulse-chase method to monitor cargo exit from the TGN. We transiently transfected the DCV cargo ANF::GFP into WT and *Eipr1* KO 832/13 cells and blocked cargo exit from the Golgi by incubating for 2 hours at 20°C (Kögel *et al*., 2013). At steady state (before the temperature block), ANF::GFP is localized more in a perinuclear region in *Eipr1* KO cells (Figure 4A), similar to insulin and mature CPE. After the temperature block (pulse), cells were returned to 37°C and incubated for different times (chase) (Figure 4B). They were then processed for GFP immunostaining and imaging. Cells were scored based on whether ANF::GFP was (1) predominantly at the TGN-region (“Golgi-like” in Figure 4C), (2) found both at the TGN-region and at the cell periphery (“Intermediate” in Figure 4C), or (3) excluded from the Golgi (“Periphery” in Figure 4C). We observed that the localization of ANF::GFP at the TGN was different between WT and *Eipr1* KO cells at all time points (Figure 4D), confirming that EIPR1 affects the distribution of DCV cargo. The distribution of ANF::GFP changed in *Eipr1* KO cells during the chase period, with most of the cells having a Golgi-like distribution at t = 0, and most of the cells having an Intermediate distribution at t = 50 min (Figure 4D). This indicates that the DCV cargo is not permanently stuck at the TGN region in *Eipr1* KO cells but is able to reach the cell periphery, although probably at a slower rate than in WT cells.

**Figure 4.**
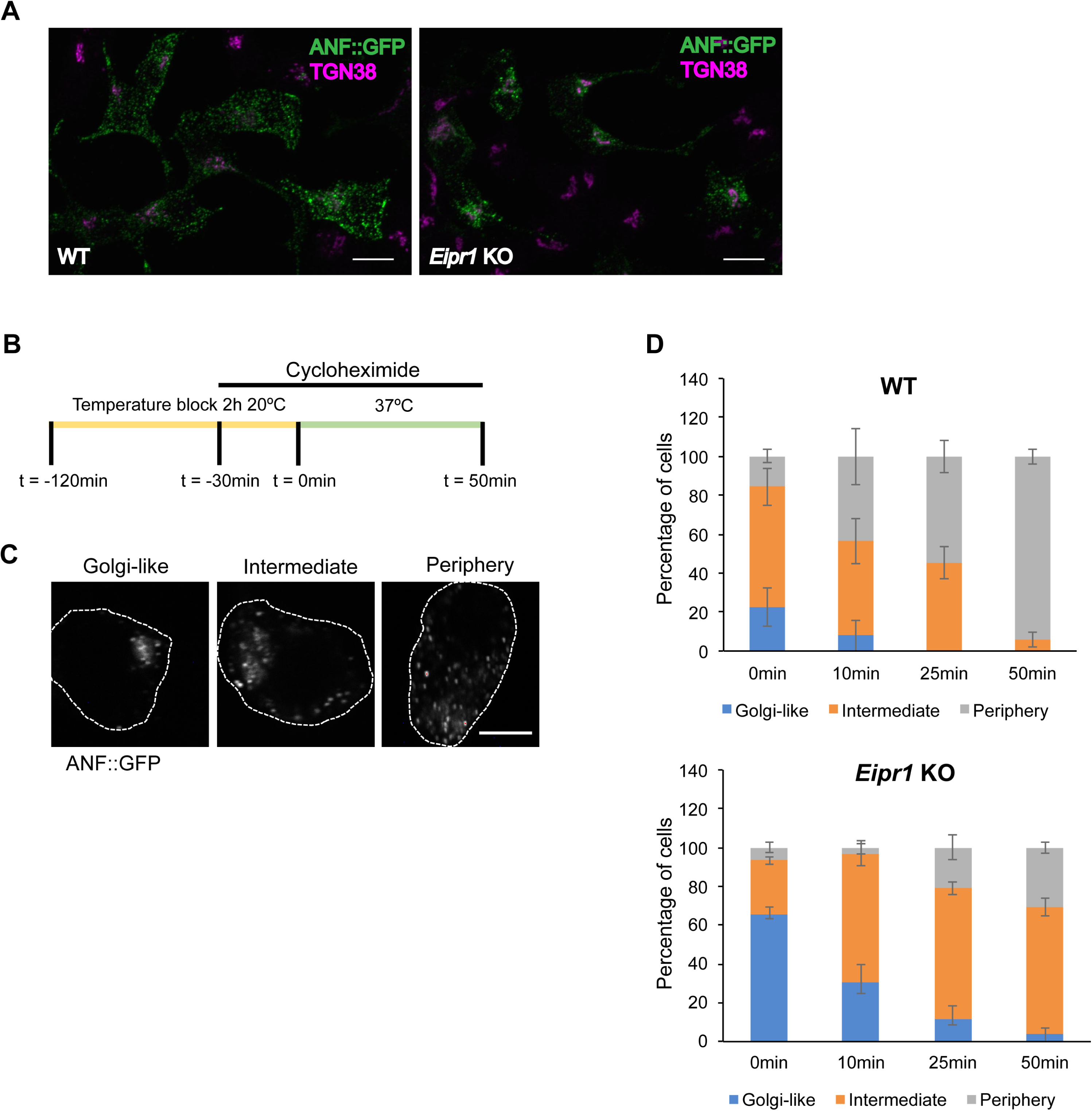
DCV cargo exit the TGN in *Eipr1* KO cells. (A) Representative images of WT and *Eipr1* KO 832/13 cells transfected with ANF::GFP and costained with anti-GFP and anti-TGN38 antibodies. In WT cells, ANF::GFP is distributed in cytoplasmic puncta, but in *Eipr1* KO cells, ANF::GFP is restricted to perinuclear puncta. Not all cells in this field of view are transfected with ANF::GFP. Scale bars: 10 μm. (B) Schematic of the pulse-chase experiment. Cells transiently transfected with ANF::GFP were incubated at 20°C for 2 h to cause the accumulation of DCV cargos at the TGN (pulse). 30 minutes before the end of the temperature block, cyclohexamide was added to block protein translation. At the end of the temperature block, cells were returned to 37°C and incubated for various times (chase) before fixation and immunostaining. (C) Representative images of the cell categories used for qualitative assessment of TGN exit: (1) ANF::GFP concentrated at the TGN region (Golgi-like), (2) ANF::GFP distributed both at the TGN and at the cell periphery (Intermediate), or (3) ANF::GFP excluded from the TGN (Periphery). Scale bar: 5 μm. (D) Percentage of WT and *Eipr1* KO cells with the indicated ANF::GFP distribution at the indicated time points. For each data point and genotype, 50 to100 cells were counted blindly. The experiment was repeated three times with similar results. The data shown for the WT are the same shown in Figure S5D of (Cattin-Ortolá, Topalidou *et al*., 2019) since these experiments were run in parallel with the same WT control.

One possible explanation for the reduced cargo in mature DCVs in the *Eipr1* KO could be that cargo is lost through the endolysosomal system. *C. elegans* EIPR-1 acts in the same genetic pathway as RAB-2 (Topalidou *et al*., 2016) and RAB-2 has been proposed to act by preventing DCV cargo from getting lost through the endolysosomal system (Edwards *et al*., 2009; Sumakovic *et al*., 2009). To test this possibility, we expressed a constitutively active form of RAB-5 (RAB-5(QL) in WT and *eipr-1* mutants in *C. elegans* to inhibit trafficking to early endosomes (Sumakovic *et al*., 2009). We found that this construct partially rescued the reduced DCV cargo levels in *eipr-1* mutant neuronal axons, similar to *rab-2* (Figure S6A,B). However, expression of rat RAB5A(QL) in *Eipr1* KO insulinoma cells did not rescue the CPE-C localization defect (Figure S6C,D). It is unclear why *C. elegans* neurons and insulinoma cells showed different effects of expressing activated RAB5.

Loss of some proteins required for DCV biogenesis (e.g. AP-3, VPS41, HID-1) leads to both loss of cargos and alterations in DCV biophysical properties such as density (Asensio *et al*., 2010, 2013; Hummer *et al*., 2017). However, we found no difference in DCV density in *Eipr1* KO cells as measured by the distribution of mature PC1/3 in membranes separated by equilibrium sedimentation through a sucrose gradient (Figure 5A,B). Thus, though levels of mature PC1/3 are reduced in *Eipr1* KO cells, the remaining PC1/3 appears to still be found in DCVs of normal density.

**Figure 5.**
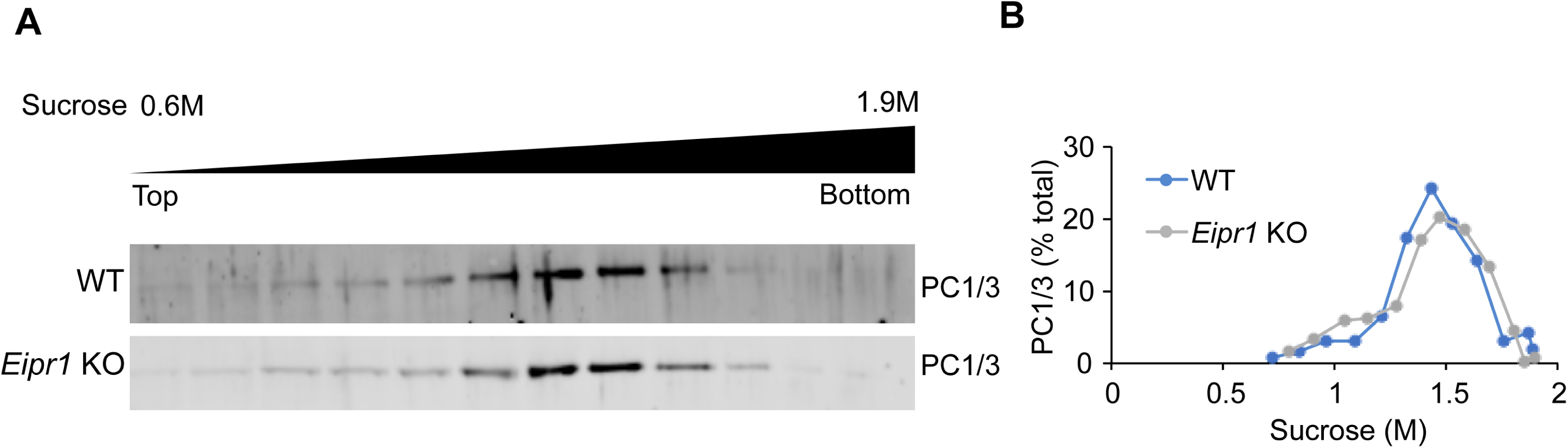
The density of DCVs is not affected in *Eipr1* KO cells. (A) Post-nuclear supernatants from WT and *Eipr1* KO cells were separated by equilibrium sedimentation through 0.6–1.9 M sucrose. Fractions were blotted with an antibody against PC1/3 (mature DCV marker). The data shown are from one representative experiment of three independent experiments with similar results. (B) Band intensity for each fraction was quantified using FIJI, presented as a percentage of total band intensity, and plotted against the sucrose concentration of the fraction.

### EIPR1 is needed for the localization of EARP subunits and their association with membranes

It was recently shown that mammalian EIPR1 interacts with the EARP and GARP complex subunits and functions with EARP in endosomal recycling and with GARP in endosome-Golgi retrograde trafficking (Gershlick *et al*., 2016; Topalidou *et al*., 2016). However, *eipr-1* mutants in *C. elegans* have behavioral and cellular phenotypes similar to EARP-specific mutants, but not GARP-specific mutants (Topalidou *et al*., 2016). EIPR1 is a WD40 domain protein, and WD40 domains often act as scaffolds for mediating protein interactions and multi-protein complex assembly (Stirnimann, Petsalaki, *et al*., 2010). To investigate whether EIPR1 is required for the localization of EARP or GARP complex subunits in insulin-secreting cells, we examined the localization of transiently transfected Myc-tagged VPS50 (the sole EARP-specific subunit), GFP and Myc-tagged VPS54 (the sole GARP-specific subunit), and Myc-tagged VPS51 and VPS53 (subunits present in both the GARP and EARP complexes). We also examined the localization of endogenous VPS50 using a commercial antibody, the only antibody we have for EARP or GARP subunits that is suitable for immunofluorescence. Unfortunately, we were unable to determine the localization of EIPR1 as we do not have an antibody that works for immunofluorescence, and tagging the *C. elegans* or rat EIPR1 proteins at either the N- or C-terminus (with either GFP or Myc) leads to a diffuse cytoplasmic localization that is probably nonphysiological. A similar issue with tagging EIPR1 was reported elsewhere (Gershlick *et al*., 2016).

In wild type cells, VPS50, VPS51, VPS53, and VPS54 all showed a punctate pattern of localization. Interestingly, the punctate localization of the EARP-specific subunit VPS50 and the EARP/GARP common subunits VPS51 and VPS53 was disrupted in *Eipr1* KO cells, with fluorescence being diffuse throughout the cytoplasm (Figure 6A,B,D, S7A-B). By contrast, the GARP-specific subunit VPS54 was still punctate and localized near the TGN in *Eipr1* KO cells (Figure 6C,D and S7C). We conclude that EIPR1 is needed for the localization of the EARP complex subunit VPS50 and the EARP/GARP common subunits, but not the GARP complex specific subunit VPS54. This result suggests that EIPR1 is preferentially required for the localization of the EARP complex.

**Figure 6.**
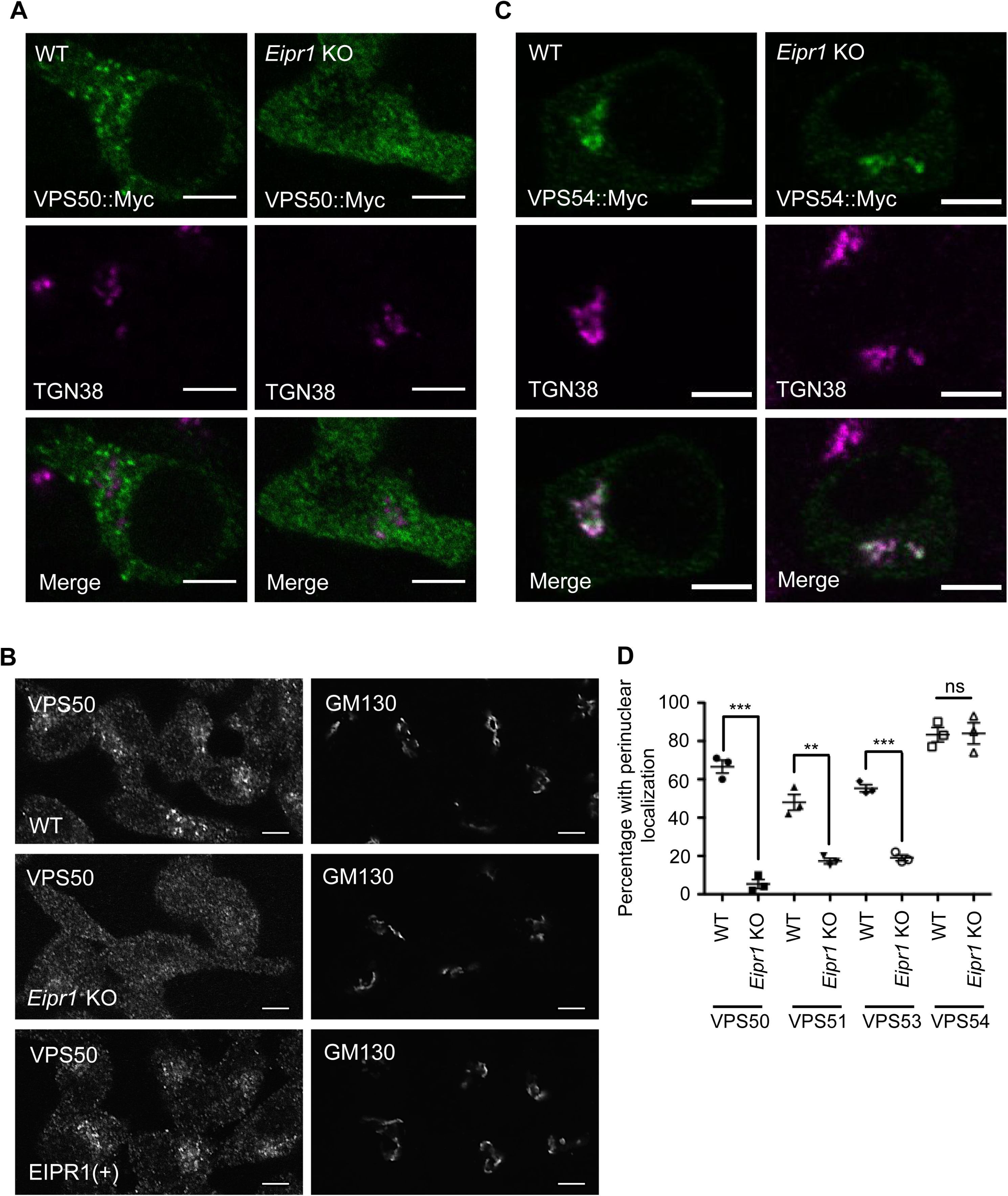
EIPR1 is required for the localization of EARP in insulin-secreting cells. (A) Representative images of 832/13 WT and *Eipr1* KO cells transfected with VPS50::13Myc (VPS50::Myc) and costained with anti-Myc and anti-TGN38 antibodies. In WT cells, VPS50::Myc is localized to puncta but in *Eipr1* KO cells fluorescence is diffuse throughout the cytoplasm. The punctate pattern of localization of VPS50::Myc overlaps only partially with TGN38. Scale bars: 5 μm. (B) Representative images of 832/13 (WT), *Eipr1* KO, and EIPR1(+) cells stained with anti-VPS50 antibody. Endogenous VPS50 is punctate in WT cells, but diffuse throughout the cytoplasm in *Eipr1* KO cells. Scale bars: 5 μm. (C) Representative images of 832/13 cells (WT) and *Eipr1* KO 832/13 (Eipr1KO) cells transfected with VPS54::13Myc (VPS54::Myc) and costained with anti-Myc and anti-TGN38 antibodies. In both WT and *Eipr1* KO cells, VPS54::Myc is localized to perinuclear puncta that largely overlap with TGN38. Scale bars: 5 μm. (D) Localization of Myc-tagged VPS50, VPS51, and VPS53, but not VPS54, is disrupted in *Eipr1* KO cells. Shown is the percentage of cells with perinuclear and punctate staining of each indicated subunit. The experiment was repeated three times and approximately 100 cells were imaged per experiment. Cells with very high expression level were not included in the counting since overexpression of the individual subunits leads to their mislocalization to the cytoplasm in WT cells. ***p<0.001, **p<0.01, ns p>0.05.

To test whether the diffuse localization of EARP subunits in the *Eipr1* KO is due to a reduced association of EARP with membranes, we fractionated 832/13 cell lysates and probed for VPS50. VPS50 was found primarily in the membrane fraction (P100) in wild-type cells, but the association of VPS50 with membranes was partially lost in *Eipr1* KO cells (Figure 7A). Similarly, VPS51 was found in the membrane fraction in wild-type cells and this association was reduced in the *Eipr1* KO (Figure 7B). By contrast, the association of the GARP-specific subunit VPS54 with membranes was not altered in *Eipr1* KO cells (Figure 7C). We conclude that EIPR1 is partially required for the proper association of EARP with membranes, but is not required for membrane-association of GARP.

**Figure 7.**
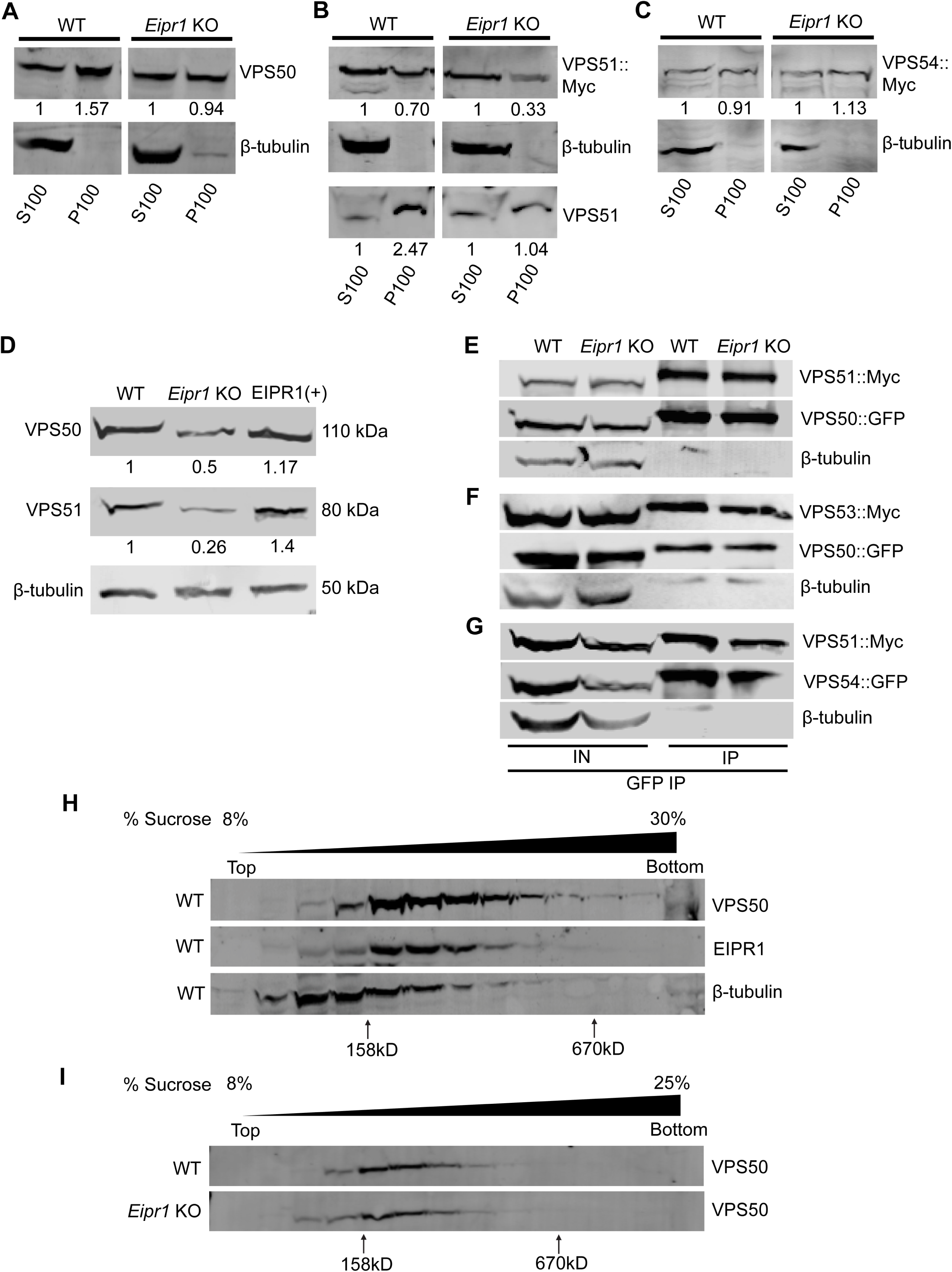
Membrane association and stability of EARP is disrupted in *Eipr1* KO insulin-secreting cells. (A) VPS50 associates with membranes in an EIPR1-dependent manner. In WT cell fractions, endogenous VPS50 was found primarily in the post-nuclear P100 membrane fraction. In *Eipr1* KO 832/13 cells, VPS50 was equally distributed between the P100 membrane fraction and the S100 cytosolic fraction. β-tubulin served as a control soluble protein. S100, P100: supernatant and pellet fractions obtained by a 100,000g spin of the cell lysate, containing cytosolic and membrane-associated proteins respectively. (B) VPS51 associates with membranes in an EIPR1-dependent manner. In WT cell fractions, VPS51::Myc was roughly equally distributed between the post-nuclear P100 membrane fraction and the S100 cytosolic fraction, and endogenous VPS51 was found primarily in the P100 membrane fraction. In *Eipr1* KO 832/13 cells (Eipr1KO), VPS51::Myc was found mostly in the soluble fraction and endogenous VPS51 was equally distributed between the membrane fraction and the cytosolic fraction. (C) VPS54 associates with membranes in an EIPR1-independent manner. In both WT and *Eipr1* KO 832/13 cell fractions, VPS54::Myc was equally distributed between the post-nuclear P100 membrane fraction and the S100 cytosolic fraction. (D) VPS50 and VPS51 levels are reduced in *Eipr1* KO cells. Protein extracts from 832/13 (WT), *Eipr1* KO 832/13 (Eipr1KO), and *Eipr1* KO 832/13 cells expressing wild type *Eipr1* (EIPR1(+)) were blotted with antibodies against VPS50 and VPS51. β-tubulin served as a loading control. (E) VPS50 interacts with VPS51 in an EIPR1-independent way. EGFP-tagged VPS50 was coexpressed with 13Myc-tagged VPS51 in 832/13 (WT) and *Eipr1* KO cells. Immunoprecipitation of VPS50::GFP pulled down VPS51::Myc independently of EIPR1, but did not pull down β-tubulin. IN: input, IP: immunoprecipitated. (F) VPS50 interacts with VPS53 in an EIPR1-independent way. EGFP-tagged VPS50 was coexpressed with 13Myc-tagged VPS53 in 832/13 (WT) and *Eipr1* KO cells. Immunoprecipitation of VPS50::GFP pulled down VPS53::Myc independently of EIPR1, but did not pull down β-tubulin. IN: input, IP: immunoprecipitated. (G) VPS51 interacts with VPS54 in an EIPR1-independent way. EGFP-tagged VPS54 was coexpressed with 13Myc-tagged VPS51 in 832/13 (WT) and *Eipr1* KO cells. Immunoprecipitation of VPS54::GFP pulled down VPS51::Myc independently of EIPR1, but did not pull down β-tubulin. IN: input, IP: immunoprecipitated (H) VPS50 and EIPR1 cofractionate on a linear 8%-30% sucrose velocity gradient. Fractions from 832/13 (WT) cell lysate were blotted with antibodies against VPS50 and EIPR1. β-tubulin served as a control soluble protein. (I) VPS50 fractionates similarly from cell lysates of 832/13 (WT) and *Eipr1* KO cells on a linear 8%-25% sucrose velocity gradient. Fractions from WT and *Eipr1* KO cell lysates were blotted with antibodies against VPS50.

### EIPR1 is not required for the physical interactions between EARP complex subunits

Because EIPR1 is needed for the proper localization of EARP complex subunits and WD40 domain proteins often serve as scaffolds for complex assembly, we examined whether EIPR1 is needed for the formation of the EARP complex. We first compared the levels of endogenous VPS50 and VPS51 in wild type and *Eipr1* KO 832/13 cells. *Eipr1* KO cells had reduced levels of VPS50 and VPS51, and this defect was rescued by expression of wild type EIPR1 (Figure 7D). Thus, EIPR1 is required for expression or stability of the individual EARP subunits. To test whether these reduced protein levels were due to reduced transcription, we examined the levels of the VPS50 and VPS51 mRNAs in WT and *Eipr1* KO cells. Quantitative PCR using total cDNA from WT and *Eipr1* KO cells showed no difference in the mRNA levels of these EARP complex subunits (Figure S2B). Thus, the EARP protein subunits are less stable in the absence of EIPR1.

To determine whether EIPR1 is required for physical interactions between the individual EARP subunits, we expressed GFP-tagged VPS50 with Myc-tagged VPS51 or Myc-tagged VPS53 in wild type and *Eipr1* KO 832/13 cells and performed coimmunoprecipitation experiments. Unlike with endogenous VPS50 and VPS51, the levels of VPS50::GFP and VPS51::Myc were not reduced in *Eipr1* KO, probably because of overexpression (Figure 7E-G, IN: input). GFP-tagged VPS50 coimmunoprecipitated with either Myc-tagged VPS51 or Myc-tagged VPS53, and these interactions were not disrupted by loss of EIPR1 (Figure 7E,F). To test whether EIPR1 is required for the interactions between GARP complex subunits, we expressed GFP-tagged VPS54 with Myc-tagged VPS51 in wild type and *Eipr1* KO 832/13 cells. GFP-tagged VPS54 coimmunoprecipitated with Myc-tagged VPS51 as expected, and this interaction was not changed in *Eipr1* KO cells (Figure 7G). These data indicate that EIPR1 is not required for interactions between the individual subunits of the EARP or GARP complexes, at least under these conditions where some subunits are overexpressed.

Finally, to further examine whether the EARP complex is disrupted in the absence of EIPR1, we subjected cell lysates from wild type and *Eipr1* KO 832/13 cells to velocity sedimentation through an 8%-30% linear sucrose gradient and blotted for VPS50. VPS50 sedimented in a broad peak between the 158 and 670 kDa standards, indicating that the protein is part of a complex (Figure 7H). EIPR1 sedimentation from the same cell lysate showed a similar peak to VPS50, suggesting that EIPR1 and VPS50 might be part of the same complex (Figure 7H). Of note, sedimentation of VPS50 was not affected by loss of EIPR1 (Figure 7I), suggesting that the EARP complex still forms in the absence of EIPR1.

### EIPR1 is required for EARP function but not GARP function

EARP is required for the recycling of cargos from recycling endosomes back to the plasma membrane (Schindler, Chen, *et al*., 2015). EIPR1 was also shown to be required for the endocytic recycling of transferrin in HAP1 cells (Gershlick *et al*., 2016). We examined whether EIPR1 is required for the recycling of transferrin in 832/13 cells and found that *Eipr1* KO 832/13 cells take up and retain slightly more transferrin than WT cells (Figure 8A), similar to results seen in HAP1 cells (Gershlick *et al*., 2016). The steady-state level and distribution of the transferrin receptor were not obviously altered in *Eipr1* KO cells (Figure 8B,C).

**Figure 8.**
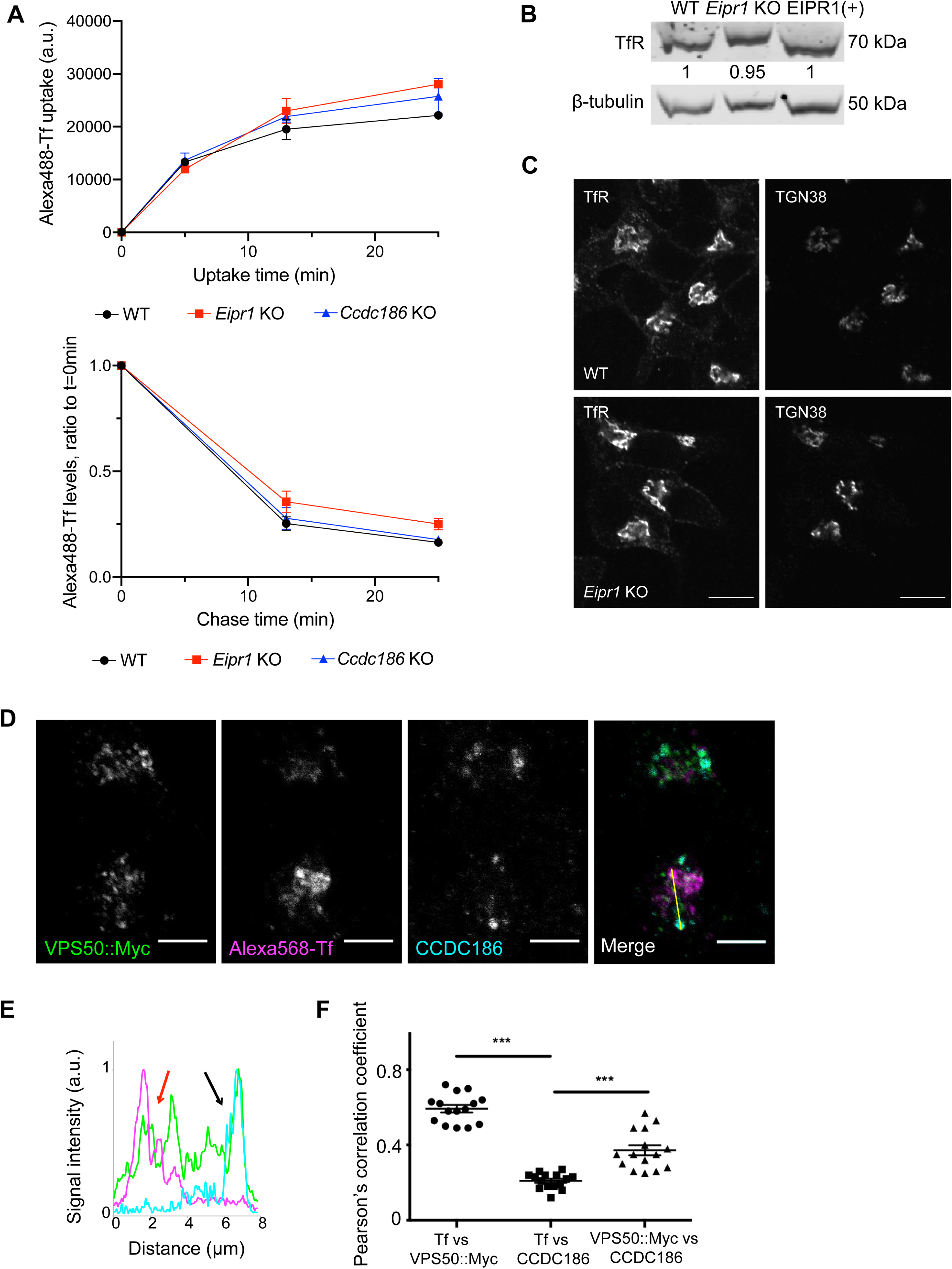
EIPR1 has a slight defect in transferrin recycling in insulinoma cells and EARP localizes to two distinct compartments. (A) EIPR1, but not CCDC186, has a slight defect in transferrin (Tf) recycling. FACS analysis of Alexa 488-labeled transferrin in WT, *Eipr1* KO, and *Ccdc186* KO cells after transferrin uptake for the indicated times (upper) and at different chase times following 25 min transferrin uptake (lower). The upper plot represents the median of Alexa 488-Tf intensity of the population of cells as a function of time. The lower plot represents the median of Alexa 488-Tf intensity of the population of cells at each time point normalized to the median intensity at t = 0 as a function of time. The experiment was repeated three times. For Tf uptake at 25 min, p=0.16, WT vs. *Eipr1* KO. For Tf recycling at 25 min, p=0.11, WT vs. *Eipr1* KO, one-way ANOVA with Bonferroni correction for multiple comparisons. (B) Levels of expression of the transferrin receptor (TfR) are not affected in *Eipr1* KO cells. Protein extracts from 832/13 (WT), *Eipr1* KO 832/13 (Eipr1KO), and *Eipr1* KO 832/13 cells expressing wild type *Eipr1* (EIPR1(+)) were blotted with an antibody against. TfR. β-tubulin served as a loading control. (C) Localization of the transferrin receptor (TfR) is not affected in *Eipr1* KO cells. Representative images of 832/13 (WT), and *Eipr1* KO (Eipr1KO) 832/13 cells costained with antibodies for endogenous TfR and TGN38. Scale bars: 5 μm. (D), (E), (F). EARP localizes to two distinct compartments: a CCDC186-positive compartment near the TGN and immature DCVs and a transferrin-positive compartment that is associated with endosomes. 832/13 cells were transiently transfected with Myc-tagged VPS50, incubated with Alexa 568-labeled transferrin (Tf) and immunostained for Myc and CCDC186. (D) Representative confocal images of cells costained for endogenous CCDC186, Alexa568-Tf and VPS50::Myc. Scale bars: 5 μm. (E) The graph shows a representative intensity plot of normalized signal intensity versus distance across the yellow line shown in (D). VPS50::Myc is shown in green, Alexa568-Tf is in magenta, and CCDC186 in cyan. Red arrow: overlap between VPS50 and Tf. Black arrow: overlap between VPS50 and CCDC186. (F) Pearson’s correlation coefficient was measured to quantify the localization between transferrin and VPS50::Myc, transferrin and CCDC186, and CCDC186 and VPS50::Myc. n=15, *** p<0.001, error bars = SEM. The experiment was repeated twice.

Depletion of GARP leads to redistribution of TGN38 to cytoplasmic vesicles thought to correspond to retrograde transport intermediates (Pérez-Victoria *et al*., 2008). We too found that siRNA knockdown of the VPS51 subunit of the GARP complex causes partial redistribution of TGN38 to cytoplasmic puncta, but the *Eipr1* KO had no obvious change in the distribution of TGN38 (Figure S8). Thus, EIPR1 is required for both the localization and function of EARP, but is not required for the localization of GARP or the function of GARP in the retrograde trafficking of TGN38.

### The EARP complex localizes to two distinct compartments

To determine whether EARP functions at distinct cellular sites to mediate endocytic recycling and DCV cargo sorting, we examined colocalization of the EARP subunit VPS50 with CCDC186 and transferrin. Transferrin is found at early and recycling endosomes and VPS50 localizes to RAB4-positive endosomes involved in transferrin receptor recycling (Schindler, Chen, *et al*., 2015). CCDC186/CCCP-1 is a coiled-coil protein that functions in DCV biogenesis in the same genetic pathway as EIPR1 and EARP (Cattin-Ortolá, Topalidou *et al*., 2019), but is not required for transferrin recycling (Figure 8A). CCDC186 is localized near immature DCVs and the TGN (Ailion *et al*., 2014; Cattin-Ortolá *et al*., 2017; Cattin-Ortolá, Topalidou *et al*., 2019), but is distinct from recycling endosomes marked by transferrin (Figure S9A). Notably, localization of CCDC186 does not require EIPR1 (Figure S9B). We observed that VPS50 is found at two distinct sites. At one site, VPS50 colocalizes with CCDC186 and at the other VPS50 colocalizes with transferrin (Figure 8D-F and S9A), consistent with EARP acting both at recycling endosomes and at the site of DCV maturation. VPS50 is found in puncta both in the perinuclear region and more peripherally and shows a localization pattern that shows little overlap with mature insulin granules in the cell periphery or the immature DCV markers syntaxin 6 and proinsulin (Figure S9C-E). By contrast, the GARP-specific VPS54 subunit was specifically localized to a perinuclear region largely overlapping TGN38 and proinsulin (Figure 6C, S9E), consistent with its role in retrograde trafficking to the Golgi as a component of the GARP complex.

## Discussion

In this study, we demonstrate that the EARP complex interacting-protein EIPR1 acts in a post-processing maturation step to regulate proper insulin secretion and distribution of mature DCV cargo. Our data indicate that cells lacking EIPR1 are capable of stimulated insulin secretion but secrete less insulin, and have less insulin and other mature DCV cargos in DCVs, supporting a role for EIPR1 in the maturation of DCVs. We further show that EIPR1 is required for localization of EARP, but not GARP, and that EIPR1 functions to control endocytic recycling and DCV cargo sorting. Consistent with its dual function, we find that the EARP complex localizes to two distinct compartments: a DCV biogenesis-related compartment and an endocytic-recycling compartment.

### EIPR1 regulates DCV cargo levels and distribution

Our studies in *C. elegans* support a role for EIPR1 and EARP in controlling DCV cargo levels. *C. elegans* mutants in *eipr-1* and the EARP complex subunits have reduced levels of cargo in mature DCVs and secrete less cargo (Paquin *et al*., 2016; Topalidou *et al*., 2016). Because these studies were based on the overexpression of exogenous DCV cargos, here we investigated the role of EIPR1 in the distribution and secretion of endogenous DCV cargo using the insulin-secreting 832/13 cell line. We found that cells lacking EIPR1 remain responsive to stimulated secretion, but secrete less insulin and contain less insulin and mature cargo in DCVs. These phenotypes are similar to those observed in neurons in *eipr-1* mutants in *C. elegans*, suggesting that neurons and endocrine cells share a conserved pathway for DCV biogenesis.

Our analysis of the levels and distribution of immature and mature DCV cargo in *Eipr1* KO cells suggests that EIPR1 acts in a post-Golgi and post-processing step to regulate DCV biogenesis. The total level of processed proprotein convertase 1/3 (PC1/3) is reduced in the absence of *Eipr1*. Also, processed and mature CPE are both reduced and misdistributed in *Eipr1* KO cells, but the level and distribution of the unprocessed form of CPE are not affected. If the reduction of PC1/3 and CPE led to a defect in the processing of proinsulin, we would expect an increase in proinsulin content in *Eipr1* KO. However, total proinsulin levels in the *Eipr1* KO were not increased, again suggesting that EIPR1 does not affect the processing of DCV cargos. Additionally, Golgi pH is unaffected in *Eipr1* KO cells. Together, these results suggest that EIPR1 is not required for early steps in the sorting and processing of DCV cargo.

Interestingly, *Eipr1* KO cells have similar defects as *Ccdc186* KO cells (Cattin-Ortolá, Topalidou *et al*., 2019), suggesting that they both act at a post-Golgi step during DCV maturation. This is in contrast to other known regulators of DCV biogenesis such as PICK1, ICA69, and HID-1 that are needed to control the budding of immature DCVs from the TGN and whose loss leads to a defect in the processing of proinsulin to insulin (Cao, Mao, *et al*., 2013; Holst, Madsen, *et al*., 2013; Du, Zhou, Zhao, Cheng, *et al*., 2016; Hummer *et al*., 2017). Our findings suggest that EIPR1 and CCDC186 act at a later step in DCV biogenesis than PICK1, ICA69, and HID-1.

### EIPR1 regulates the localization of the EARP complex and its association with membranes

The WD40 domain protein EIPR1 was identified as an interactor of the EARP and GARP complexes in rat insulin-secreting cells and human neuroglioma cells (Gershlick *et al*., 2016; Topalidou *et al*., 2016). Additionally, two mass spectrometry interactome data sets found that EIPR1 interacts with EARP subunits in human HEK293T or HeLa cells (Hein *et al*., 2015; Huttlin *et al*., 2015). VPS50 was shown to pull down VPS51, VPS52, VPS53, and EIPR1 as a stoichiometric complex (Hein *et al*., 2015), indicating that EIPR1 may form a stable complex with EARP. Although WD40 domain proteins often act as scaffolds for the assembly of large protein complexes (Stirnimann, Petsalaki, *et al*., 2010), our data suggest that EIPR1 is not needed for the formation of the EARP or GARP complex. First, EIPR1 is not required for interactions between individual subunits of the EARP and GARP complexes, as shown by coimmunoprecipitation experiments. Second, sedimentation of VPS50 was not affected by loss of EIPR1, suggesting that the EARP complex still forms in the absence of EIPR1. By contrast, we find that EIPR1 is needed for the stability of the individual EARP subunits, the localization of the EARP complex subunits, and association of EARP with membranes, supporting the model that EIPR1 recruits the EARP complex to its site of action and stabilizes it there.

Although localization of the EARP subunits is disrupted in the absence of EIPR1, localization of the GARP-specific VPS54 subunit is not affected. A recent study also showed that VPS54::GFP stably expressed in H4 cells is localized at the TGN region in both wild type and *Eipr1* knockdown (KD) cells, but FRAP analysis showed that EIPR1 contributes to GARP recruitment to the TGN (Gershlick *et al*., 2016), suggesting that EIPR1 is not needed for localization of GARP to the TGN per se, but is required to recruit GARP efficiently.

### EIPR1 participates in EARP-specific functions

EARP and EIPR1 were recently shown to participate in the endocytic recycling of transferrin (Schindler, Chen, *et al*., 2015; Gershlick *et al*., 2016). We also found that EIPR1 is needed for normal endocytic recycling of transferrin in insulinoma cells. To test whether EIPR1 might also function with GARP, we examined the distribution of TGN38 in *Eipr1* KO and *Vps51* KD cells. In the absence of GARP, TGN38 is partially redistributed to cytoplasmic vesicles (Pérez-Victoria *et al*., 2008), but loss of EIPR1 had no obvious effect on the distribution of TGN38. Additionally, *C*. *elegans* GARP mutants were shown to have enlarged lysosomes in coelomocytes, but *eipr-1* and *vps-50* mutants do not have enlarged lysosomes (Topalidou *et al*., 2016). These results suggest that EIPR1 does not participate in GARP-specific functions, which agrees with the model that EIPR1 is required for the localization of EARP but not GARP. However, EIPR1 was reported to be required for the retrograde traffic of the Shiga toxin B subunit (STxB), whose trafficking also depends on GARP (Gershlick *et al*., 2016). We were unable to examine the trafficking of STxB in insulinoma 832/13 cells since we found that these cells do not take up STxB. These data together suggest that if EIPR1 functions with GARP, it is required for only a subset of GARP functions.

### EARP localizes to two distinct compartments

The dual functionality of EIPR1 and EARP in endocytic recycling and DCV cargo retention prompted us to examine the cellular distribution of EARP. We found that EARP localizes to two distinct cellular sites, one that is associated with endosomes and one associated with CCDC186 near the TGN and immature DCVs. The endosomal recycling and DCV cargo sorting functions of EARP and EIPR1 may reflect two independent functions or may be interconnected. It is possible that the EARP complex moves from endosomes to the Golgi to participate in the retrieval of DCV cargo or sorting factors from an endosomal compartment. Alternatively, a pool of EARP localized near the TGN and immature DCVs may function in DCV biogenesis independently of a second pool of EARP that acts to traffic cargo out of endosomes.

### A connection between endosomal trafficking and DCV biogenesis?

We have demonstrated here and elsewhere that EARP, an endosomal-recycling complex, and the EARP-interacting protein EIPR1 are involved in DCV cargo retention (Topalidou *et al*., 2016), raising the possibility that DCV maturation at or near the TGN may require input from endosomal compartments. Several other studies have also suggested that there may be a role of endosomes in DCV biogenesis or maturation (Klumperman *et al*., 1998; Vo *et al*., 2004; Edwards *et al*., 2009; Sumakovic *et al*., 2009; Bäck *et al*., 2010; Topalidou *et al*., 2016; Zhang *et al*., 2017).

There are several possible ways that DCV maturation may be connected to endosomes. Studies in *C. elegans* suggested that RAB-2 acts in DCV biogenesis by ensuring that cargo is retained in mature DCVs and is not lost to the endolysosomal system (Edwards *et al*., 2009; Sumakovic *et al*., 2009). EIPR-1 and RAB-2 act in the same genetic pathway (Topalidou *et al*., 2016), suggesting that EIPR1 might also act to retain cargos in mature DCVs and prevent their loss to the endolysosomal system. In agreement with this model, we found that disrupting trafficking to the endolysosomal system by overexpressing a constitutively active form of RAB-5 partially restored levels of DCV cargo in *C. elegans eipr-1* mutants. However, expression of an analogous activated RAB5A construct did not restore the CPE localization defect in *Eipr1* KO insulinoma cells, so the precise relationship between DCV maturation and endolysosomal trafficking remains unclear.

A second possible connection of endosomes to DCV maturation is the removal of cargos from immature DCVs. The AP-1 adaptor that is involved in trafficking between the trans-Golgi and endosomes has been shown to associate with immature DCVs and to mediate the removal of syntaxin 6 and mannose 6-phosphate receptors from immature DCVs (Dittie *et al*., 1996; Klumperman *et al*., 1998). Once removed, such proteins possibly follow the endosomal route (Feng and Arvan, 2003; Arvan and Halban, 2004). In support of this idea, we found that EIPR1 is needed for the removal of carboxypeptidase D (CPD) from immature DCVs (Cattin-Ortolá, Topalidou *et al*., 2019). CPD is localized to the TGN and immature DCVs, but is mostly absent from mature DCVs (Varlamov *et al*., 1999). In the absence of EIPR1, CPD remains in mature DCVs (Cattin-Ortolá, Topalidou *et al*., 2019), suggesting that EIPR1 ensures the removal of CPD from immature DCVs. The RAB-2 effector CCDC186 was also shown to act in a similar way (Cattin-Ortolá, Topalidou *et al*., 2019). We propose that EIPR1 and CCDC186 play a role in ensuring that the proper amount and type of cargo remains in mature DCVs.

Another possible connection of endosomes to DCV maturation is the recycling and retrieval of DCV cargos from the plasma membrane. Following DCV exocytosis, transmembrane DCV cargos may be recycled back to nascent DCVs via an endosomal pathway (Vo *et al*., 2004; Bäck *et al*., 2010). Additionally, a recent study identified a possible role for the secretory cell-specific Munc13-4 paralog BAIAP3 in this DCV recycling pathway (Zhang *et al*., 2017). BAIAP3 was shown to localize to late and recycling endosomes and to be needed for DCV maturation and for more general endosome recycling to the TGN (Zhang *et al*., 2017). Because EARP and EIPR1 act in recycling plasma membrane proteins like transferrin out of endosomes, perhaps EARP and EIPR1 are also needed for the trafficking of recycled DCV membrane cargo out of endosomes to the cellular compartments where DCVs are formed and mature. It will be interesting to determine whether DCV membrane cargos are recycled from the plasma membrane through endosomes in an EIPR1 and EARP-dependent manner.

## Materials & methods

### Cell culture

The 832/13 cell line is an INS-1-derived clone isolated by the lab of Dr. Christopher Newgard (Duke University School of Medicine) (Hohmeier, Hubner, *et al*., 2000) and obtained by Dr. Duk-Su Koh via Dr. Ian Sweet (University of Washington). Cell lines were grown in RPMI 1640-GlutaMAX^™^ (GIBCO) medium supplemented with 10% FBS (RMBIO), 1 mM sodium pyruvate (GIBCO), 10 mM HEPES (GIBCO), 1X Pen/Strep (GIBCO), and 0.0005% 2-beta-mercaptoethanol at 5% CO_2_ and 37°C. Cells were transfected using Lipofectamine 2000 (Thermo Fisher) according to the manufacturer’s instructions.

### Constructs

The plasmids VPS50::13Myc, VPS51::13Myc, VPS53::13Myc, VPS54::13Myc, VPS54::GFP, and VPS50::GFP were a gift from Juan Bonifacino (Pérez-Victoria *et al*., 2008; Pérez-Victoria and Bonifacino, 2009; Schindler, Chen, *et al*., 2015). The ANF::GFP plasmid (rat atrial natriuretic factor fused to GFP) was described (Hummer *et al*., 2017). The EIPR1_pBabe-hygro construct (pET222) used for making EIPR1(+) stable lines was constructed by amplifying rat EIPR1 cDNA from an 832/13 cDNA library using primers:

oET513: 5’-ccatggatccatggaagacgacgccccg-3’ and
oET514: 5’-ctgagaattctcagagcagtatgtggtacttcagtgc-3’

The PCR product was digested by EcoRI/BamHI and cloned into EcoRI/BamHI-digested pBabe-hygro (a gift from Suzanne Hoppins).

The RAB5A[Q79L]-pEGFP-N1 construct (pET259) used for expressing mutant RAB5A in 832/13 cells was made by PCR amplifying rat RAB5A using primers:

oJC388: 5’-gatctcgagctcaagcttcgATGGCTAATCGAGGAGCAACAAGACC-3’
oJC389: 5’-ccatggtggcgaccggtggatcGTTACTACAACACTGACTCCTGGCTG-3’

The PCR product was inserted in pEGFP-N1 vector using Gibson cloning (Gibson *et al*., 2009) and mutagenized using the Quickchange II site directed mutagenesis kit (Agilent #200523) and the following primer set:

oET645: 5’-AATATGGGATACAGCTGGCCtAGAACGGTATCATAGCCTA-3’
oJC388: 5’-TAGGCTATGATACCGTTCTaGGCCAGCTGTATCCCATATT-3’

### *Eipr1* knock out using CRISPR editing

To knock out EIPR1, we performed Cas9-mediated genome editing via homology-directed repair (HDR) in 832/13 cells using the protocol described (Ran, Hsu, *et al*., 2013).

For designing guide RNAs we used the online CRISPR design tool (Ran, Hsu, *et al*., 2013) and selected three guide RNAs that recognize sequences in or near the first exon of rat *Eipr1*:

guide 1: 5’-gacgacgccccggtgatctacggg-3’
guide 2: 5’-gagcccgagtcccgcctcaccagg-3’
guide 3: 5’-gtatcatggaagacgacgccccgg-3’

The guide RNAs were cloned into pSpCas9(BB)-2A-GFP vector using the indicated protocol (Ran, Hsu, *et al*., 2013). The efficiency of the cloned guide RNAs was tested using the SURVEYOR nuclease assay (Figure S1B) according to the manufacturer’s instructions (Surveyor Mutation Detection kit, Transgenomic). Guide RNA #1 (plasmid pET45) was selected for all subsequent experiments.

To design the homology-directed repair (HDR) template we used the pPUR (Clontech) vector as a backbone and cloned approximately 1.5 kb *Eipr1* homology arms upstream and downstream of the puromycin selection cassette (Figure 1A and S1A). The HDR template was constructed using Gibson assembly (plasmid pET69).

To cotransfect the CRISPR plasmid (carrying Cas9 and the guide RNA) and the HDR template, cells were grown in two 10-cm petri dishes to near confluency. Cells were cοtransfected with 7 μg CRISPR plasmid and 7 μg non-linearized HDR template using Lipofectamine 3000 according to the instructions (Thermo Fisher). 48 hours after transfection, the media was removed and replaced with new media together with 1 μg/ml puromycin. The puromycin selection was kept until individual clones could be picked, grown in individual dishes, and tested for CRISPR editing.

Individual puromycin-resistant clones were tested for proper CRISPR editing of the *Eipr1* gene by extracting genomic DNA and performing PCR (Figure S1C). The primers used for the PCR screening of positive clones were the following:

oET236: 5’-gaggtccgttcacccacag-3’ (hybridizes just upstream of the left homology arm).
oET237: 5’-gcctggggactttccacac-3’ (hybridizes in the SV40 promoter that drives the expression of the puromycin resistance gene).

5 out of 16 puromycin-resistant clones showed the band indicative of insertion of the puromycin cassette into *Eipr1*. To test for homozygosity of the insertion, we performed PCR using primers that amplify the wild-type *Eipr1* locus:

oET236: 5’-gaggtccgttcacccacag-3’ (hybridizes just upstream of the left homology arm).
oET200: 5’-gagcagtatccaacacacacctc-3’ (hybridizes just downstream of the Cas9 cleavage site and in the first *Eipr1* intron).

Of the 16 clones tested, three did not show the wild-type band with primers oET236 and oET200 (Figure S1C) and were subsequently tested for EIPR1 expression by Western blot. Clone #3 showed no EIPR1 expression by Western (Figure 1B) and was kept as the *Eipr1* KO line.

### Lentiviral production, infection of cells, and selection of stable lines

Platinum-E (PlatE) retroviral packaging cells (a gift from Suzanne Hoppins) were grown for a couple of generations in DMEM-GlutaMAX^™^ (GIBCO) medium supplemented with 10% FBS (RMBIO), 1X Pen/Strep (GIBCO), 1 μg/ml puromycin, and 10 μg/ml blastocidin at 5% CO_2_ and 37°C. On day one, approximately 3.6 × 10^5^ PlatE cells per well were plated in a six-well dish in DMEM-GlutaMAX^™^ medium supplemented with 10% FBS and 1X Pen/Strep. On day two, a mix of 152 μl Opti-MEM (Thermo Fisher), 3 μg EIPR1_pBabe-hygro DNA, and 9 μl Fugene HD transfection reagent (Promega) was incubated for 10 minutes at room temperature and transfected into each well. On day three, the media was removed and replaced with new PlatE media. On day four, approximately 1.5 × 10^5^ *Eipr1* KO 832/13 cells per well were plated in a six-well dish in RPMI 1640-GlutaMAX, supplemented with 10% FBS, 1 mM sodium pyruvate, 10 mM HEPES, 1X Pen/Strep, and 0.0005% 2-beta-mercaptoethanol. 3 μl of 8 mg/ml hexadimethrine bromide (Sigma) was added to each well. The supernatant of the PlatE cells (48 hours viral supernatant) was collected with a sterile syringe, passed through a 0.45 micron filter, and added to the *Eipr1* KO cells. The *Eipr1* KO cells were incubated for 5-8 hours at 5% CO_2_ and 37°C, then the media was changed and replaced with new media. The cells were incubated overnight at 5% CO_2_ and 37°C. On day five, the supernatant was removed from the *Eipr1* KO cells and replaced with the supernatant from the PlatE cells (72 hours viral supernatant) after passing through a 0.45 micron filter. 3 μl of 8 mg/ml hexadimethrine bromide was added in each well and the cells were incubated for 5-8 hours. The media was replaced with new RPMI 1640-GlutaMAX media. On day six, the *Eipr1* KO cells were collected, transferred to a 10-cm petri dish, and 200 μg/ml hygromycin was added. The cells were grown under hygromycin selection until individual clones could be picked and tested for EIPR1 expression.

### Insulin and proinsulin secretion

Cells were grown in 24-well plates to near confluency, washed twice with PBS, and incubated for 1 hour in 200 μl per well resting buffer (5 mM KCl, 120 mM NaCl, 24 mM NaHCO_3_, 1 mM MgCl_2_, 15 mM HEPES pH 7.4). The medium was collected, cleared by centrifugation, and stored at −80^°^C. The cells were incubated for 1 hour in 200 μl per well stimulating buffer (55 mM KCl, 25 mM glucose, 70 mM NaCl, 24 mM NaHCO_3_, 1 mM MgCl_2_, 2 mM CaCl_2_, 15 mM HEPES pH 7.4). After stimulation, the medium was cleared by centrifugation and stored at −80°C. The cells were washed once with PBS, harvested in PBS, and extracted in 100 μl per well acid-ethanol solution (absolute ethanol:H_2_0:HCl, 150:47:3). The pH of the acid-ethanol solution was neutralized by addition of 20 μl of 1M Tris Base per 100 μl of acid ethanol and the samples were stored at −80°C.

Samples were assayed for insulin or proinsulin content using ELISA according to the instructions of the manufacturers (Rat/Mouse insulin ELISA, Millipore, #EZRMI-13K. Rat/Mouse proinsulin ELISA, Mercodia, #10-1232-01). Secreted insulin and proinsulin levels were normalized against total cellular protein concentration and were presented as fraction of the wild type under stimulating conditions (Fig 1C,D left panels). These values were then normalized against total cellular insulin or proinsulin levels (Fig 1C,D middle panels) to give the secretion data normalized against total insulin or proinsulin content (Fig 1C,D right panels).

### Quantitative RT-PCR

WT and *Eipr1* KO cells were grown in 10 cm plates, harvested in 1 ml of TRIzol (Invitrogen), and frozen at −80°. Total RNA was isolated following the manufacturer’s protocol and cDNA was synthesized from 1 μg RNA using the QuantiTekt Reverse Transcription kit (Qiagen) according to the manufacturer’s instructions. Each 10 μl qPCR reaction contained 1 μl of cDNA and 5 μl of 2× Sybr Green Master Mix (Kappa Biosystems). Absorbance was measured over 40 cycles using a CFX Connect Real-Time System (Biorad). The cycle quantification value (Cq, cycle at which the amplification curve crosses a prescribed threshold value) for each sample was measured using the provided software and normalized to actin control. The primers used were the following:

Proinsulin 1 F: 5’-atggccctgtggatgc-3’
Proinsulin 1 R: 5’-tcagttgcagtagttctccagttg-3’
PC1/3 F: 5’-atgaagcaaagaggttggactc-3’
PC1/3 R: 5’-ttaattcttctcattcagaatgtcc-3’
Actin F: 5’-atggatgacgatatcgctgc-3’
Actin R: 5’-ctagaagcatttgcggtgc-3’
VPS50 F: 5’-atgcaaaaaatcaaatctcttatgacccgg-3’
VPS50 R: 5’-tcgtttaggtctgtctatatcatctatagc-3’
VPS51 F: 5’-atggcggccgcggcagctgtggggcctggc-3’
VPS51 R: 5’-gccgcgctcgcagatgacctcgacaacact-3’

### Exocytosis assay

832/13 cells stably expressing NPY-pHluorin were transfected (FuGene, Promega) with NPY-mCherry. At 1 d after transfection, cells were washed once with PBS, dislodged using media, and transferred onto poly-L-lysine coated 22 mm glass coverslips. After an additional 2 d, cells were reset for 2 hrs in low K+ Krebs-Ringer buffer with 1.5 mM glucose, washed once with low K+ Krebs-Ringer buffer with 1.5 mM glucose, and coverslips were transferred to an open imaging chamber (Life Technologies). Cells were imaged close to the coverslips, focusing on the plasma membrane (determined by the presence of NPY-mCherry-positive plasma membrane docked vesicles), using a custom-built Nikon spinning disk confocal micrscope at a resolution of 512 × 512 pixels. Images were collected for 100 ms at 10 Hz at room temperature with a 63X objective (Oil Plan Apo NA 1.49) and an ImageEM X2 EM-CCD camera (Hamamatsu, Japan). Following baseline data collection (15 s), an equal volume of Krebs-Ringer buffer containing 110 mM KCl and 30.4 mM glucose was added to stimulate secretion and cells were imaged for an additional 80 s. At the end of the experiment, cells were incubated with Krebs-Ringer buffer containing 50 mM NH_4_Cl, pH 7.4, to reveal total fluorescence and to confirm that the imaged cells were indeed transfected. Movies were acquired in MicroManager (UCSF) and exported as tiff files. Movies were analyzed using ImageJ software by counting events and measuring cell area.

### Constitutive secretion assay

WT and *Eipr1* KO cells were seeded on 12-well microtiter plates. When they reached subconfluency they were transfected using Lipofectamine 2000 with a plasmid expressing GFP fused to a signal peptide at its N-terminus (ssGFP) (Hummer *et al*., 2017). 48h after the transfection, cells were incubated with resting media (114 mM NaCl, 4.7 mM KCl, 1.2 mM KH_2_PO_4_, 1.16 mM MgSO_4_, 20 mM HEPES, 2.5 mM CaCl_2_, 25.5 mM NaHCO_3_, 3 mM glucose), for 1h at 37°C. The secretion media was collected, centrifuged at 20,000g for 10 minutes at 4°C to remove cells and cell debris, and the supernatant was collected. The pelleted cells were lysed on ice in lysis buffer (50 mM Tris-HCl, pH 8.0, 150 mM NaCl, 1% Triton X-100, 1 mM EDTA, protease inhibitor cocktail without EDTA (Pierce)). The cell lysate was centrifuged at 20,000g at 4°C for 10 minutes and the supernatant was collected. Protein concentration was measured using the BCA assay. GFP fluorescence of the media (secreted GFP) and the lysate were measured using a plate reader (Excitation = 485 nm, Emission = 525 nm, cutoff = 515 nm). Background fluorescence was measured from the media and lysate from non-transfected cells.

### Immunostaining

Approximately 1-2 × 10^5^ cells per well were plated onto cover slips (Thomas Scientific #121N79) placed in 24-well cell culture plates. Cells were transfected with Lipofectamine 2000 according to manufacturer’s instruction at least 24h after seeding. After 24 to 48 hours, the cells were rinsed twice with PBS and fixed with 4% paraformaldehyde (made in PBS) for 20 minutes at room temperature. The cells were rinsed twice with PBS and permeabilized with 0.5% Triton X-100 in PBS for 5 minutes at room temperature. The cells were rinsed twice with PBS and placed in 5% milk in PBS for 1 hour at room temperature. Cells were stained with primary antibodies in 0.5% milk in PBS at room temperature for 1 hour. The following primary antibodies were used: mouse monoclonal anti-c-Myc (1:1000, Santa Cruz, sc-40), rabbit polyclonal anti-CCDC132/VPS50 (1:50, Sigma #HPA026679), mouse monoclonal anti-GFP (1:200 to 1:350, Santa Cruz, #sc-9996), mouse monoclonal anti-insulin (1:350, Sigma, #K36AC10), mouse monoclonal anti-proinsulin (1:100, Abcam, #ab8301), rabbit polyclonal anti-TGN38 (1:350, Sigma, #T9826), rabbit polyclonal anti-pro CPE (1:100), rabbit polyclonal anti-CPE-C terminus (1:100) (Fricker *et al*., 1996), rabbit polyclonal anti CPE-N terminus (1:100) (Fricker *et al*., 1990), mouse monoclonal anti-transferrin receptor (1:100, ThermoFisher, #13-6800), mouse monoclonal anti-syntaxin 6 (1:100, Abcam, #ab12370), rabbit polyclonal anti-CCDC186 (1:150, Novus #NBP1-90440). The cells were then washed with PBS three times for 5 minutes each, and incubated with rhodamine anti-rabbit secondary antibody (1:1000, Jackson Immunoresearch #111-025-144), Alexa Fluor 488 anti-rabbit secondary antibody (1:1000, Jackson Immunoresearch, #115-545-152), Alexa Fluor 488 anti-mouse secondary antibody (1:1000, Jackson Immunoresearch, #115-545-146), and Rhodamine Ret-X anti mouse secondary antibody (1:1000, Jackson Immunoresearch #715-295-150) at room temperature for 1 hour. The cells were washed with PBS three times for 5 min each, mounted onto glass slides using Vectashield (Vector laboratories H1000) or Prolong Diamond (Life Technologies P36965) and examined by fluorescence microscopy. Images were obtained using a Nikon 80i wide-field compound microscope with a 60X oil objective (numerical aperture = 1.4) or an Olympus FLUOVIEW FV1200 confocal microscope with a 60X UPlanSApo oil objective (numerical aperture = 1.35). The acquisition software used for the Nikon was NIH elements and for the Olympus it was Fluoview v4.2. Pearson’s correlation coefficients were determined using Fiji and the coloc-2 plugin by taking maximum intensity projections of z-stacks and drawing a line around each individual cell. For the quantification of the insulin or CPE that is retained at the Golgi, z-stack images were obtained using a Nikon 80i wide-field compound microscope. Maximum intensity projections were quantified using Fiji. The final fluorescence intensity is either the total fluorescence in a region of interest that includes the Golgi divided by either the fluorescence of a region of the same size in the cytoplasm of the cell (Golgi/Cytoplasm) or by the background fluorescence of a region of the same size outside the cell (Golgi/Background). We were not able to examine the localization of EIPR1 by immunostaining since our antibody does not work in immunofluorescence.

### Protein extraction

For protein extraction or coimmunoprecipitation, approximately 4 × 10^6^ 832/13 cells were plated onto 10-cm plates. When transfection was required, cells were transfected 24 hours later with 15 ug of the relevant plasmid (or 8 μg each if two plasmids were transfected). After 24 to 48 hours the cells were washed with cold PBS twice and harvested in lysis buffer containing 50 mM Tris, pH 7.5, 150 mM NaCl, 1% NP40 and protease inhibitor cocktail (Pierce). Lysates were transferred to microcentrifuge tubes and passed 10 times through a 20G needle followed by incubation for 30 min at 4°. Lysates were centrifuged at 20,000g for 15 min at 4°.

### Immunoblotting

For the blots shown in Figures 1B, 7 D-G, and 8B, membranes were blocked in 3% milk in TBST (50 mM Tris pH 7.4, 150 mM NaCl, 0.1% Tween 20) for 1 hour at room temperature and stained with the relevant antibodies in 3% milk in TBST overnight, followed by three 5-minute washes in TBST. For the blots shown in Figure 3A-D, 5A, 7A-C, H, and I, membranes were blocked with Odyssey® Blocking Buffer (PBS, 927-10100). Antibodies were incubated in the same buffer and washed with PBST (137 mM NaCl, 2.7 mM KCl, 10mM Na_2_HPO_4_, 1.8mM KH_2_PO_4_, pH 7.4 supplemented with 0.1% Tween-20). The following primary antibodies were used: rabbit polyclonal anti-GFP (1:1000, a gift from Dr. Alexey Merz), mouse monoclonal anti-c-Myc (1:1000, Santa Cruz, sc-40), mouse monoclonal anti-beta-tubulin (1:1000, ThermoFisher, BT7R, #MA5-16308), mouse monoclonal anti-beta-tubulin (1:1,000, DHSB, E7), rabbit polyclonal anti-PCSK1 (PC1/3) (1:1000, Sigma, #SAB1100415), rabbit polyclonal anti-TSSC1/EIPR1 (1:1000, Thermo Scientific, #PA5-22360), rabbit polyclonal anti-CCDC132/VPS50 (1:1000, Sigma, #HPA026679), rabbit polyclonal anti-VPS51 (1:1000, Atlas antibodies, #HPA061447), rabbit polyclonal PC2 (1:1000, #13/4, a gift from Sharon Tooze (Dittié and Tooze, 1995), rabbit polyclonal anti-pro-CPE (1:1000, a gift from Dr. Lloyd D. Fricker; antiserum to pro-CPE was generated to the peptide QEPGAPAAGMRRC coupled to maleimide-activated keyhole limpet hemocyanin (KLH); this peptide corresponds to the 12 residues of mouse/rat/human pro-CPE with an added Cys on the C-terminus (for coupling to the carrier protein, KLH)), rabbit polyclonal anti-CPE-C (1:1000, antiserum was raised against the 9-residue peptide KMMSETLNF corresponding to the C-terminus of mouse CPE (Fricker *et al*., 1996)), rabbit polyclonal anti-CPE-N (1:1000, antiserum was raised to the N-terminal 15 amino acids of bovine CPE (Fricker *et al*., 1990)), mouse monoclonal anti-transferrin receptor (1:1000, ThermoFisher, #13-6800), Membranes were stained with the relevant secondary antibodies in 3% milk in TBST, followed by three 5-minute washes in TBST. The secondary antibodies used were an Alexa Fluor 680-conjugated goat anti-mouse antibody (1:20,000, Jackson Laboratory, #115-625-166), Alexa Fluor 790-conjugated donkey anti-mouse antibody (1:20,000, Jackson Laboratory, #715-655-150), or Alexa Fluor 680-conjugated goat anti-rabbit antibody (1:20,000, Jackson Laboratory, #115-625-144). A LI-COR processor was used to develop images.

### pH measurement of the late-Golgi compartment

WT and *Eipr1* KO cells stably expressing St6Gal1::pHluorin were generated as follows. HEK293T cells were maintained in DMEM with 10% fetal bovine serum under 5% CO2 at 37°C. Lentivirus was produced by transfecting HEK293T cells with FUGW, psPAX2, and pVSVG using Fugene HD according to the manufacturer’s instructions. Two days after transfection, the medium was collected and filtered (0.45 µm). Medium containing the lentivirus was then applied to 832/13 cells in suspension at varying ratios. Transduction efficiency was screened using an epifluorescence microscope. The protocol to measure the pH of the late-Golgi compartment (Hummer *et al*., 2017) was adapted to a plate-reader format. An equal number of WT and *Eipr1* KO 832/13 cells stably expressing St6Gal1::pHluorin were plated on clear-bottom black-wall 96-well plates and grown to confluence. Cells were washed once with Tyrode buffer (119 mM NaCl, 2.5 mM KCl, 2 mM CaCl_2_, 2 mM MgCl_2_, 25 mM HEPES, 30 mM glucose, pH 7.4) and incubated for 10 minutes either in Tyrode buffer of in KCl enriched buffer at different pHs (125 mM KCl, 20 mM NaCl, 0.5 mM CaCl_2_, 0.5 mM MgCl_2_, and 25 mM MES (at pH 5.5 or 6) or 25 mM HEPES (at pH 6.5, 7, 7.5, 8, or 8.5). The KCl-enriched buffer was supplemented with 5 μM nigericin (Sigma Aldrich) and 5 nM monensin (Sigma Aldrich). The fluorescence of each well was measured using a Varioskan Lux plate reader (Thermo Scientific) (Excitation=485 nm, Emission=520 nm). For each buffer, cell type, and independent biological replicate, the reading was repeated three times. Calibration curves were generated for each cell type and each independent repetition. The absolute pH values were extrapolated from the calibration curve using the fluorescence of the cells incubated with Tyrode buffer.

### ANF-GFP pulse-chase

To monitor the exit of DCV cargo from the TGN, we used a protocol similar to the one described (Kögel *et al*., 2013)). WT and *Eipr1* KO 832/13 cells were seeded on glass coverslips and grown for 24 hours. We then transfected 100 ng of ANF::GFP (Hummer *et al*., 2017) with Lipofectamine 2000 for 12-16 hours at 37°C in complete growth medium. Cells were then incubated at 20°C in PBS for 2 h in a conventional incubator (pulse) to block the formation of DCVs. 30 minutes before the end of the low temperature block, 10 μg/ml of cyclohexamide was added to the PBS to block the synthesis of new ANF::GFP. The PBS was then exchanged for growth medium and the cells were shifted to 37°C (chase) for the indicated times and then were fixed with 4% paraformaldehyde (PFA), stained, and imaged as described (see immunostaining section). Cells were scored in three categories: those that had most of the ANF::GFP concentrated at the TGN (“Golgi-like”), those that had ANF::GFP both at the TGN-region and at the cell periphery (“Intermediate”), and those where the ANF::GFP was excluded from the TGN (“Periphery”). 50 to 100 cells per time point were counted for each genotype. The experimenter was blind to the genotypes of the cell lines used and the time point. The experiment was repeated three times with similar results.

### Worm strains

Worm strains were cultured using standard methods (Brenner, 1974). The following strains were used in this study:

KG1395 *nuIs183*[*Punc-129*::NLP-21-Venus, *Pmyo-2*::NLS-GFP] III
XZ1026 *rab-2(nu415)* I; *nuIs183*[*Punc-129*::NLP-21-Venus, *Pmyo-2*::NLS-GFP] III
XZ1055 *eipr-1(tm4790)* I; *nuIs183*[*Punc-129*::NLP-21-Venus, *Pmyo-2*::NLS-GFP] III
XZ1996 *yakIs30*[*Prab-3*::mCherry::RAB-5(QL), *Pmyo-2*::mCherry]
XZ2021 *rab-2(nu415)* I; *nuIs183*[*Punc-129*::NLP-21-Venus, *Pmyo-2*::NLS-GFP] III; *yakIs30*[*Prab-3*::mCherry::RAB-5(QL), *Pmyo-2*::mCherry]
XZ2022 *eipr-1(tm4790)* I; *nuIs183*[*Punc-129*::NLP-21-Venus, *Pmyo-2*::NLS-GFP] III; *yakIs30*[*Prab-3*::mCherry::RAB-5(QL); *Pmyo-2*::mCherry]

### Worm imaging and image analysis

Young adult worms were washed off plates with M9 and centrifuged for 3 min at 2,000 rpm. 5 μl of the worm pellet was placed on 2% agarose pads together with 5 μl of 100 mM sodium azide and the animals were anesthetized for 10 min. The dorsal cord of the animals around the vulva was imaged in all worms using a Nikon 80i wide-field compound microscope. Maximum intensity projections of z-stacks were quantified using ImageJ. The final fluorescence intensity for each animal is the averaged total fluorescence in five regions of interest minus the background fluorescence of a region of the same size next to the dorsal cord.

### Equilibrium sedimentation

Wild type and *Eipr1* KO 832/13 cells were grown in 15-cm tissue culture plates until confluent. Cells were washed twice with ice-cold PBS, transferred to 15 ml conical tubes, and centrifuged at 300 g for 10 minutes at 4°C. Cells were washed once with 5 ml of SH buffer (10 mM HEPES pH 7.2, 0.3 M sucrose, 1 mM PMSF and protease inhibitor cocktail), resuspended in 1 ml SH buffer, passed once through a 21-gauge needle, and homogenized using a ball bearing device (Isobiotec, 18 μm clearance). The post-nuclear lysate was collected after centrifugation at 1000 g for 8 minutes. A 0.6-1.9 M continuous sucrose gradient was prepared in 10 mM HEPES pH 7.2 in ultracentrifuge tubes (Beckman Coulter) that were coated with Sigmacote (Sigma #SL2). The post-nuclear supernatant was loaded onto the gradient and centrifuged at 30,000 rpm in an SW41 rotor for 14-16 hours at 4°C. Fractions of 750 μl each were collected from top to bottom. The sucrose concentration for each fraction was determined by measuring the Brix using a Misco palm abbe digital refractometer. Fractions were analyzed by immunoblotting using an antibody to PC1/3 (Sigma, #SAB1100415). Band intensity was quantified using FIJI and plotted against the sucrose concentration of the fraction.

### Cell fractionation

We used a method similar to the one described (Cattin-Ortolá *et al*., 2017). Specifically, WT 832/13 or *Eipr1* KO 832/13 cells were seeded on a 24-well cell culture plate and grown until sub-confluence. Where indicated, cells were transfected with VPS51::13Myc or VPS54::13Myc for 48 hours before fractionation using Lipofectamine 2000, according to the manufacturer’s instructions. Cells were washed twice with ice-cold PBS, transferred into microcentrifuge tubes in PBS, and centrifuged at 500g for 10 minutes at 4°C. Cells were resuspended in lysis buffer (20 mM HEPES pH 7.4, 250 mM sucrose supplemented with protease inhibitors from Pierce) and disrupted by repeated passage through a 30-gauge needle. Cell lysates were centrifuged twice at 1,000g for 10 minutes at 4°C. The post-nuclear supernatant was transferred into an ultracentrifuge tube (Beckman-Coulter #343775) and centrifuged at 100,0000g for 1h at 4°C in a TLA100 rotor. The supernatant was removed and supplemented with 6X SDS loading dye. The pellet was washed once with lysis buffer, resuspended in an equal volume of lysis buffer, and supplemented with SDS loading dye. Samples were analyzed by immunoblotting as described above.

### Coimmunoprecipitation

The anti-GFP nanobody was expressed and purified as described (Topalidou *et al*., 2016). Approximately 1/10^th^ of the supernatant (see protein extraction) was kept as input control for subsequent immunoblot analysis. The remaining lysate was incubated with 20 μg anti-GFP nanobody bound to magnetic beads for two to four hours at 4°C. The beads were washed three times with lysis buffer and resuspended in Laemmli loading buffer. The input and immunoprecipitated samples were resolved on 8% SDS-polyacrylamide gels and blotted onto PVDF membranes.

### Cofractionation on velocity sucrose gradients

WT or *Eipr1* KO 832/13 cells were grown to confluence in 15-cm tissue culture plates. Cells were washed twice with ice-cold PBS, transferred into microcentrifuge tubes in PBS, and centrifuged at 500g for 10 minutes at 4°C. Cells were resuspended in lysis buffer (50 mM Tris-Cl pH 7.6, 150 mM NaCl, 1% NP-40, 1 mM EDTA, protease inhibitor cocktail from Pierce), kept on ice for 20 minutes and centrifuged at 20,000g for 10 minutes at 4°C. Continuous sucrose gradients were prepared by overlaying 8% sucrose on 30% sucrose (or 8%-25% as indicated) in 50 mM Tris-Cl pH 7.6, 150 mM NaCl, 1% NP-40, 1 mM EDTA, in a 3.5 ml ultracentrifuge tube (Beckman Coulter #349622). The tubes were sealed with parafilm, set horizontally for 2 hours at 4°C, and then vertically for 1 hour at 4°C. The clarified lysate (or sizing standards) was loaded on top of the gradient. Tubes were centrifuged at 100,000g for 16 h at 4°C in a SW50 rotor. Fractions were collected from top to bottom and supplemented with 6X SDS sample loading buffer. EARP complex subunits were analyzed by immunoblotting using antibodies to the endogenous proteins. Sizing standards (Bio-Rad #151-1901) were analyzed by Coomassie-stained SDS-PAGE gels.

### Transferrin uptake and recycling assays and immunostaining assays

For the transferrin uptake and recycling assays, WT, *Eipr1* KO, and *Ccdc186* KO 832/13 cells were seeded onto 24-well plates and grown in complete media until they reached confluence. For the uptake, cells were placed in warm uptake medium (Serum free RPMI + 1% BSA + 25 mM HEPES + 50 µg/ml Alexa 488-transferrin (Invitrogen #T13342)) for the indicated times (Figure 8A Top). For the recycling, cells were placed in warm uptake medium for a 25-minute pulse. Medium was then exchanged for complete RPMI medium and cells were chased for the indicated times.

Cells were then transferred to ice, washed twice with ice-cold PBS, and detached on ice with 10mM EDTA in PBS for 30 minutes to 1 hour with manual agitation and gentle pipetting. Detached single cells were transferred into a microcentrifuge tube and fixed at 4°C with 4% PFA (final concentration ∼ 3.5%) for 10 minutes on a Nutator to avoid clumping. Fixed cells were washed with PBS and analyzed by FACS using a LSRII (BD Biosciences). Data were analyzed using FlowJo.

For transferrin immunostaining, cells were seeded onto cover slips in 24-well plates for 24 to 48 hours. Cells were then placed in warm uptake medium for a 25-minute pulse (Serum free RPMI + 1% BSA + 25 mM HEPES + 50 µg/ml Alexa 568-transferrin (Invitrogen #T23365)), washed twice with PBS, and fixed with 4% PFA as already described.

### VPS51 knockdown by RNAi

VPS51 knockdown was performed using stealth siRNA from Life Technologies (# 10620312-353281 D10, 5’-GAUGGACAGUGAGAcGGACAUGGUG-3’). Transfection of oligonucleotides (20 nM) was done using Lipofectamine 2000 as follows: cells were seeded on cover slips placed in 24-well cell culture plates until they reached approximately 50% confluence. On days one and three, cells were transfected with 1 μl of oligonucleotides and 1 μl Lipofectamine according to the manufacturer’s instructions. On day five, cells were stained with the relevant antibodies following standard immunostaining procedure.

### Statistics

Data were tested for normality by a Shapiro-Wilk test. When the data did not pass the normality test, we used the Kruskal-Wallis test followed by Dunn’s test to investigate whether there was statistical significance between groups of three or more. When data passed the normality test, we used a 1-way ANOVA test with Bonferroni correction when making comparisons among groups of three or more, and an unpaired t-test for comparisons between groups of two.

## Abbreviations

DCV: dense-core vesicle
EARP: endosome-associated recycling protein
GARP: Golgi-associated retrograde protein
TGN: trans-Golgi network
PC1/3: proprotein convertase 1/3
CPE: carboxypeptidase E
CPD: carboxypeptidase D

## Acknowledgements

We thank Christopher Newgard, Ian Sweet and Duk-Su Koh for the 832/13 cell line, with the support of the UW DRC Cell Function and Analysis Core; Suzanne Hoppins for the pBabe vector, platE cells, and protocol for lentiviral production and infection; Juan Bonifacino for the EARP and GARP subunit plasmids; Alex Merz and Sharon Tooze for antibodies; Lloyd Fricker for antibodies and helpful discussions. JC was supported in part by an NIH Institutional Training Grant for Neurobiology (T32 GM007108). This work was supported by an American Diabetes Association grant #1-17-JDF-064 and by NIH grant R01 GM124035 to CSA, and by a University of Washington Diabetes Research Center Pilot and Feasibility Award (NIH grant P30 DK017047), an Ellison Medical Foundation New Scholar Award, and by NIH grants R00 MH082109 and R01 GM121481 to MA. The authors declare no competing financial interests.

## Supplementary Figures

**Figure S1.**
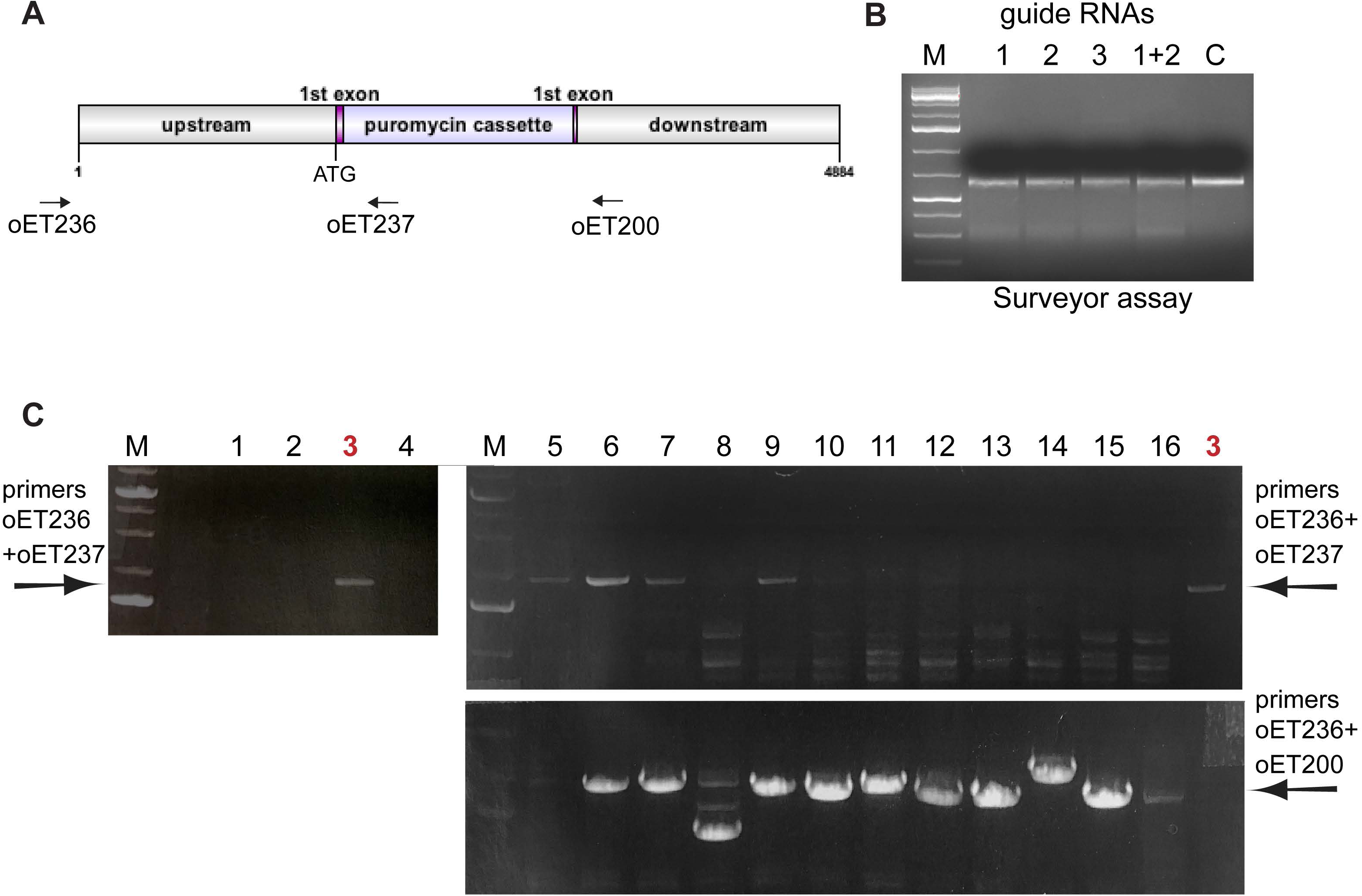
*Eipr1* KO strategy using the CRISPR technology. (A) To make the *Eipr1* KO 832/13 cell line, Cas9-induced DNA cleavage was used to insert a puromycin cassette in the first exon of *Eipr1*. oET236, oET237 and oET200 are the primers used in (C) for detecting the positive clones. (B) Surveyor nuclease assay testing the efficiency of three different guide RNAs (1, 2, 3) and the combination of guide RNAs #1 and #2 (1+2). All three guide RNAs recognize sequences at or around the 1^st^ exon of rat *Eipr1*. Guide RNA #1 was used for all subsequent experiments. C: control. (C) PCR detection of the *Eipr1* CRISPR positive clones using the indicated primers. Primers oET236 and oET237 detect clones positive for the puromycin insertion. Primers oET236 and oET200 detect clones that contain the wild type product. Clones #3 and #5 were selected as candidate *Eipr1* KOs. A Western blot showed that clone #3 lacked EIPR1 expression (Figure 1A), but that #5 still expressed the wild type EIPR1 product. Clone #3 was used for all the *Eipr1* KO experiments in this study.

**Figure S2.**
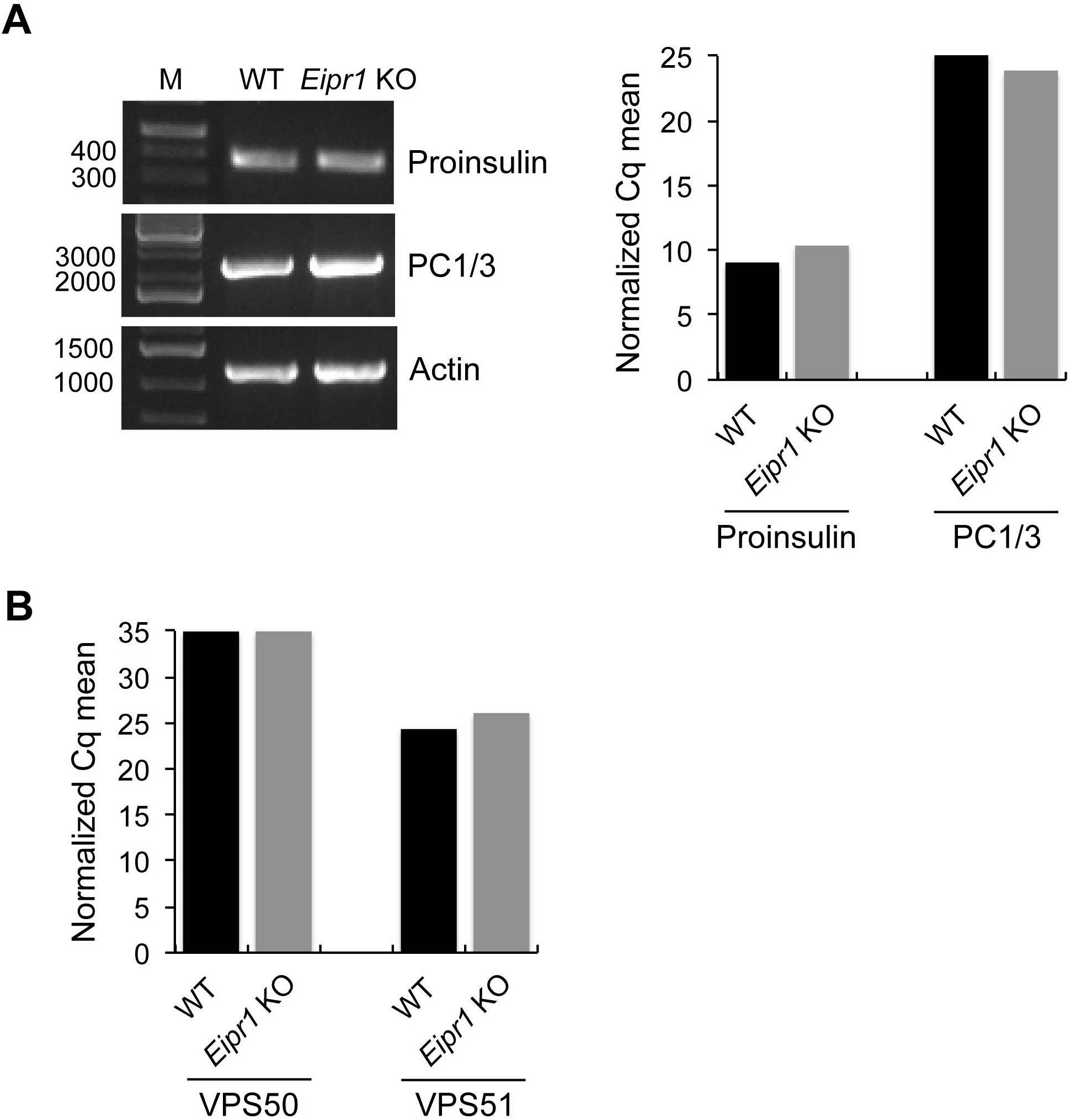
EIPR1 knock out does not affect transcription of DCV cargos and EARP or GARP subunits. No change in transcript abundance of (A) proinsulin and PC1/3 or (B) VPS50 and VPS51 in *Eipr1* KO cells as measured by qRT-PCR. Actin served as an internal control. The Cq mean for the target genes was normalized against the Cq mean for the actin control.

**Figure S3.**
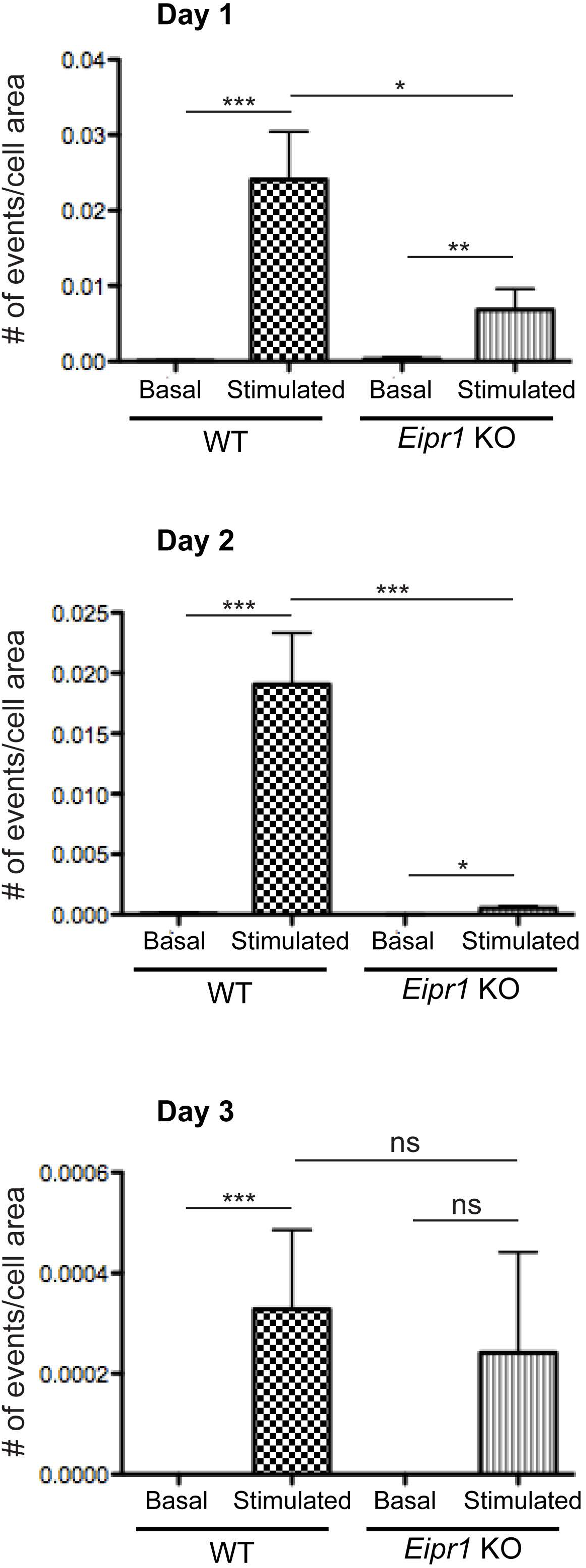
*Eipr1* knock out cells have exocytosis defects. WT and *Eipr1* KO 832/13 cells stably expressing NPY-pHluorin were imaged using spinning-disk confocal microscopy. Cells were reset for 2 hrs in low K+ Krebs-Ringer buffer with 1.5 mM glucose. Cells were then imaged in low K+ Krebs-Ringer buffer with 1.5 mM glucose for 15 s, and stimulated with 60 mM K+ and 16.7 mM glucose for 80 s. Images (100 ms exposure) were collected at 10 Hz. Exocytotic events were hand-counted. Bar graphs show the number of exocytotic events per second normalized to cell surface area. ***, p<0.001, \**p*<0.05, ns p>0.05, error bars = SEM. Three experiments were performed on different days and plotted separately. The *Eipr1* KO showed reduced stimulated exocytosis on days 1 and 2, but no significant difference from WT on day 3. However, very few events were observed on day 3, even for WT (note the scale of the y-axis), suggesting problems on that day with the stimulation protocol or ability to detect events. On day 1, n=47 cells for WT (four coverslips imaged separately with 20, 12, 7, and 8 cells); n=43 cells for *Eipr1* KO (three coverslips with 17, 14, and 12 cells). On day 2, n=79 cells for WT (four coverslips with 26, 13, 22, and 18 cells); n=61 cells for *Eipr1* KO (three coverslips with 29, 24, and 8 cells). On day 3, n=30 cells for WT (two coverslips with 12 and 18 cells); n=11 cells for *Eipr1* KO (two coverslips with 8 and 3 cells).

**Figure S4.**
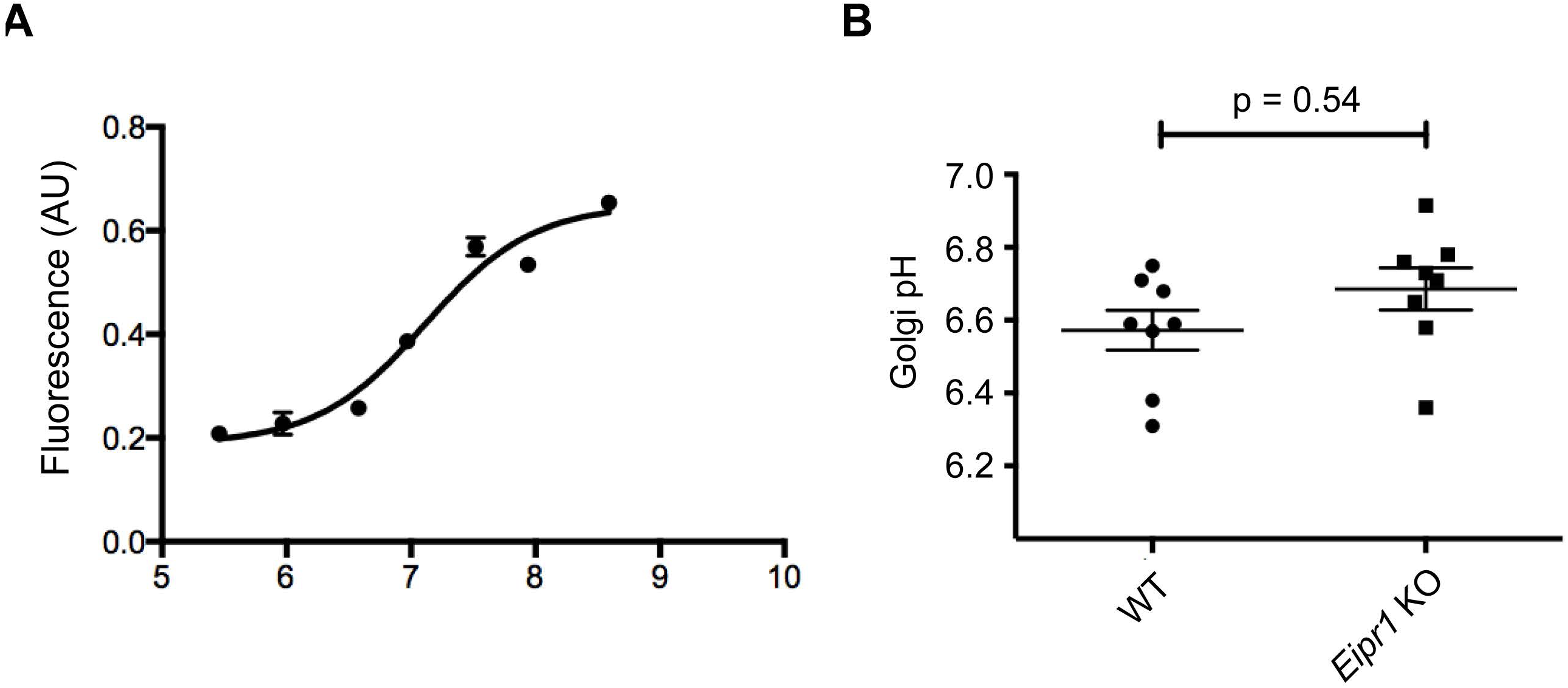
The pH of the Golgi is not significantly changed in *Eipr1* KO cells. (A) Example calibration curve for Golgi pH measurements based on measuring the fluorescence of TGN-targeted pHluorin in solutions of defined pH. For each biological replicate of WT and *Eipr1* KO 832/13 cells tested for pH, an individual calibration curve such as the one shown was obtained from WT and *Eipr1* KO cells grown in the same 96-well plate as the test samples and exposed to buffers of decreasing pH (8.5–5.5) in the presence of nigericin and monensin (see Materials and Methods). Each data point shows the mean of the fluorescent measurements (in arbitrary units (AU)) from cells in three different wells. Error bars = SEM. (B) The late-Golgi compartment is not more acidic in *Eipr1* KO 832/13 cells. The fluorescence of TGN-targeted pHluorin was measured in WT and *Eipr1* KO cells. The absolute pH value of each sample was extrapolated from a paired calibration curve (as in A). The data show mean ± SEM; n = 8 for WT and *Eipr1* KO. The data shown for the WT are the same shown in Figure S3 of (Cattin-Ortolá, Topalidou *et al*., 2019) since these experiments were run in parallel with the same WT control.

**Figure S5.**
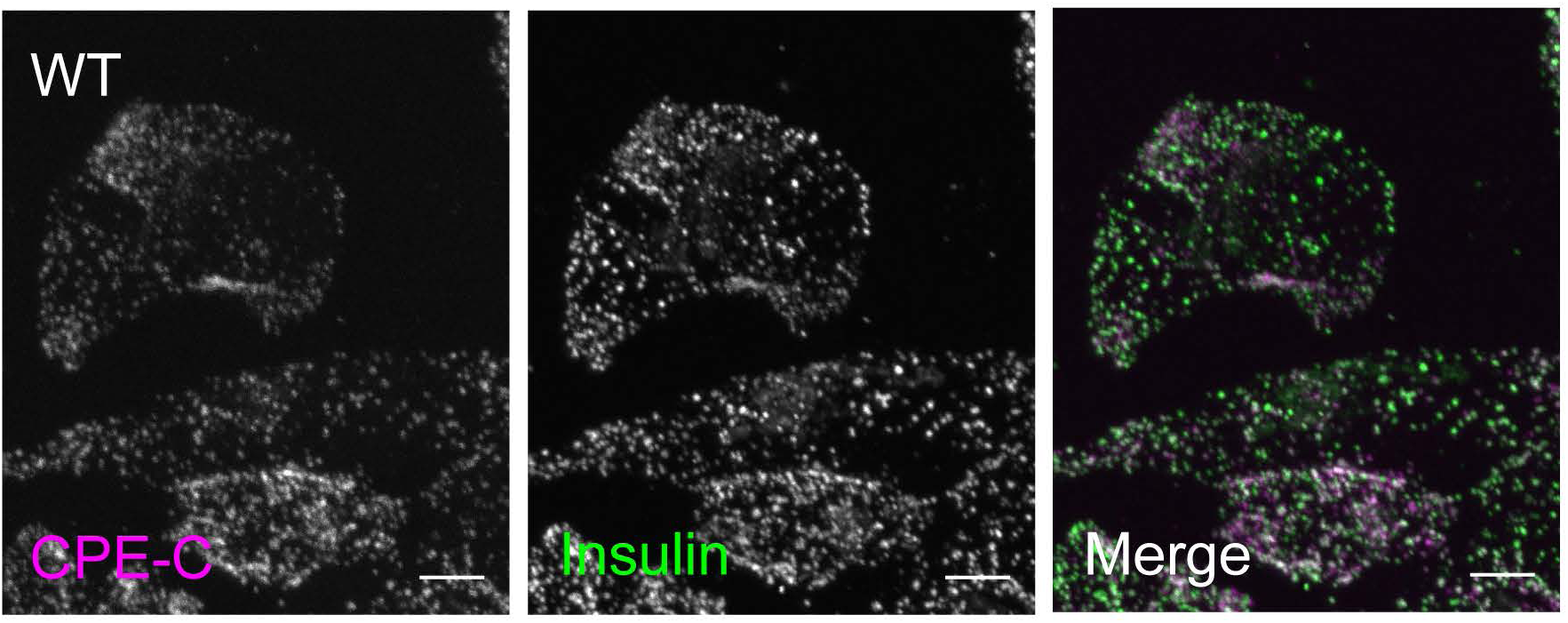
Mature CPE is localized in insulin-containing DCVs. Representative images of 832/13 (WT) cells costained with antibodies for mature CPE (CPE-C) and insulin. Scale bars: 5 μm.

**Figure S6.**
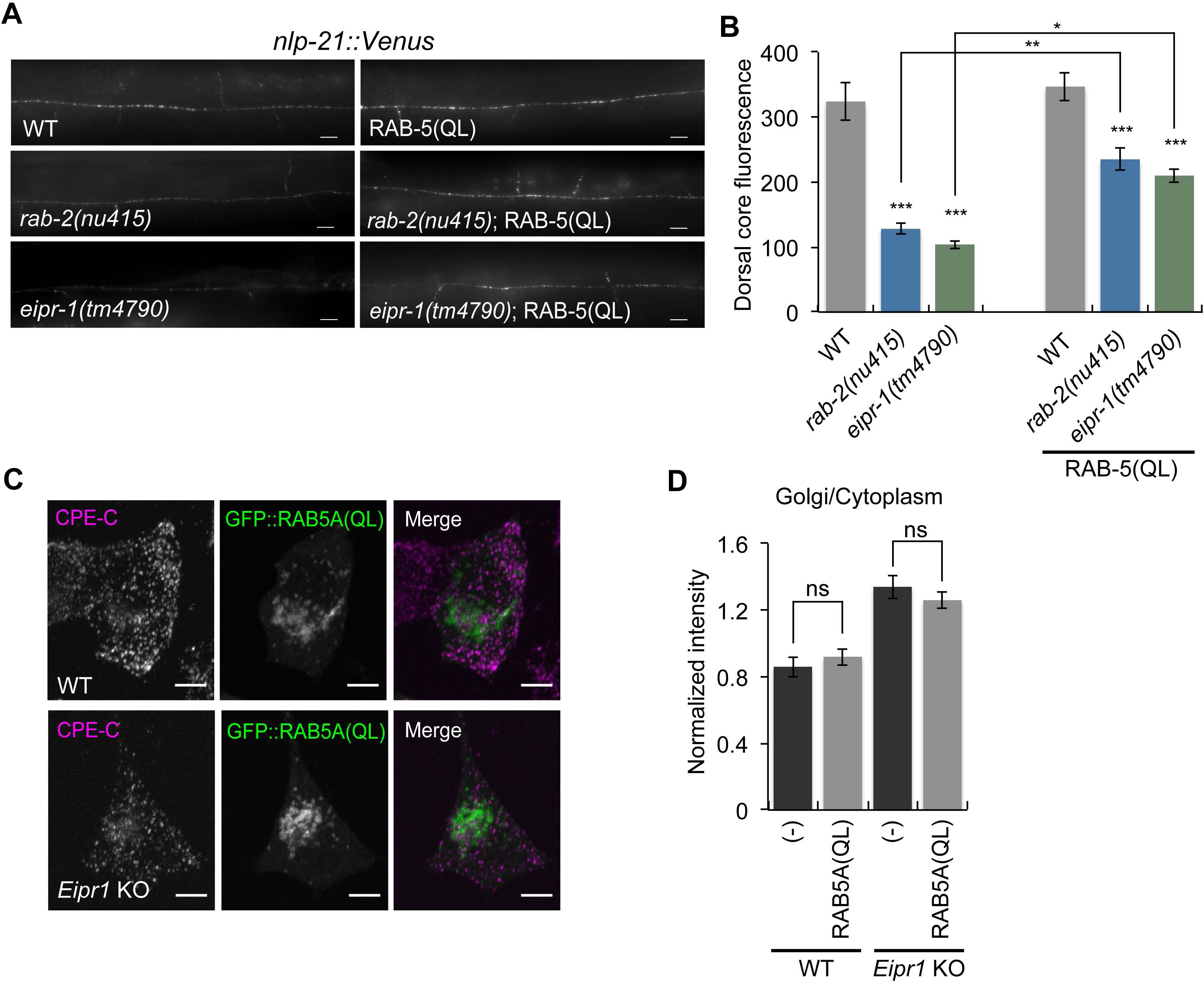
Expression of constitutively active RAB-5(QL) in neuronal cells partially rescues the NLP-21::Venus defect of *eipr-1 C. elegans* mutants. (A) Representative images of NLP-21::Venus fluorescence in motor neuron axons of wild type (WT), *rab-2(nu415),* and *eipr-1(tm4790)* mutant strains, with or without RAB-5(QL). Scale bar: 10 μm. (B) Quantification of NLP-21::Venus fluorescence levels in motor neuron axons of the indicated strains. The mean fluorescence intensity is given in arbitrary units. *rab-2(nu415)* and *eipr-1(tm4790)* mutants have decreased levels of NLP-21::Venus fluorescence. This phenotype is partially rescued by expressing RAB-5(QL). (n = 10-20, error bars = SEM, * p<0.05, ** p<0.01, *** p<0.001). (C) Representative images of of 832/13 (WT) and *Eipr1* KO cells transfected with GFP::RAB5A(QL) and costained with antibodies for GFP and mature CPE (CPE-C). Scale bars: 5 μm. (D) *Eipr1* KO cells have an increased Golgi/cytoplasmic ratio of mature CPE relative to wild type cells and this phenotype is not rescued by expression of GFP::RAB5A(QL). Fluorescence of a region of interest that includes the TGN divided by the fluorescence of a region of the same size in the cytoplasm, in WT and *Eipr1* KO with or without GFP::RAB5A(QL). (n=11-16, error bars = SEM, ns p>0.05)

**Figure S7.**
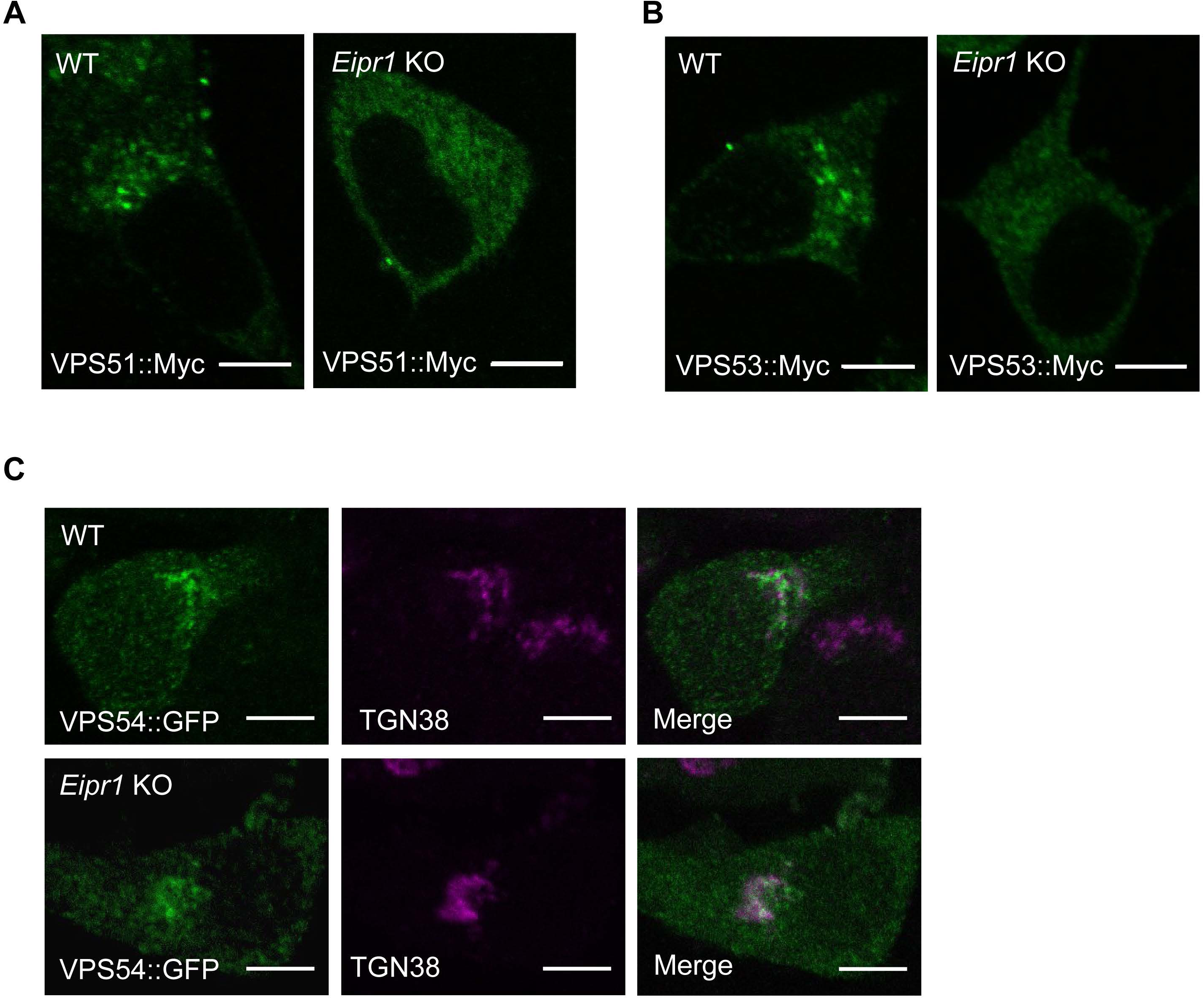
Localization of VPS51::Myc and VPS53::Myc is disrupted in *Eipr1* KO cells, but VPS54::GFP is not affected. (A) Representative images of 832/13 (WT) and *Eipr1* KO 832/13 (Eipr1KO) cells transfected with VPS51::13Myc (VPS51::Myc) and stained with anti-Myc antibody. VPS51::Myc is punctate in WT cells, but diffuse throughout the cytoplasm in *Eipr1* KO cells. Scale bars: 5 μm. (B) Representative images of 832/13 (WT) and *Eipr1* KO 832/13 (Eipr1KO) cells transfected with VPS53::13Myc (VPS53::Myc) and stained with anti-Myc antibody. VPS53::Myc is punctate in WT cells, but diffuse throughout the cytoplasm in *Eipr1* KO cells. Scale bars: 5 μm. (C) Representative images of 832/13 (WT) and *Eipr1* KO 832/13 (Eipr1KO) cells transfected with VPS54::GFP (VPS54::GFP) and stained with anti-GFP and TGN38 antibodies. In both WT and *Eipr1* KO cells, VPS54::GFP is localized to perinuclear puncta that largely overlap with TGN38. Scale bars: 5 μm.

**Figure S8.**
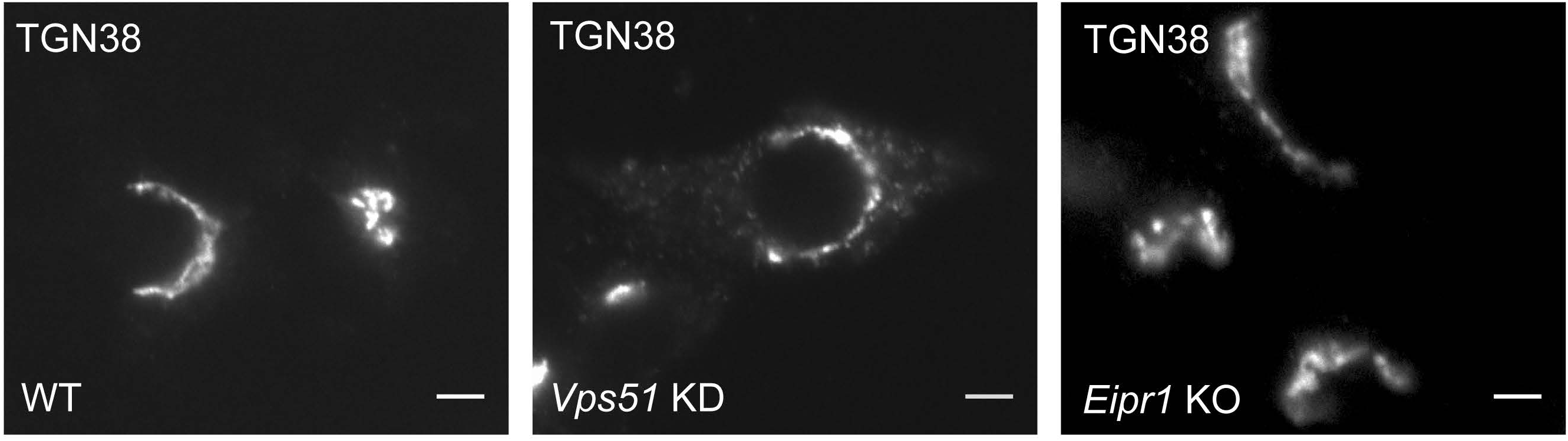
TGN38 is redistributed in *Vps51* knockdown but not *Eipr1* KO cells. Representative images of 832/13 cells (WT), *Vps51* siRNA knockdown 832/13 cells (Vps51KD), and *Eipr1* KO cells (Eipr1KO) stained for TGN38. TGN38 is partially redistributed to cytoplasmic puncta in VPS51 knockdown cells, but is still localized to the Golgi in *Eipr1* KO cells.

**Figure S9.**
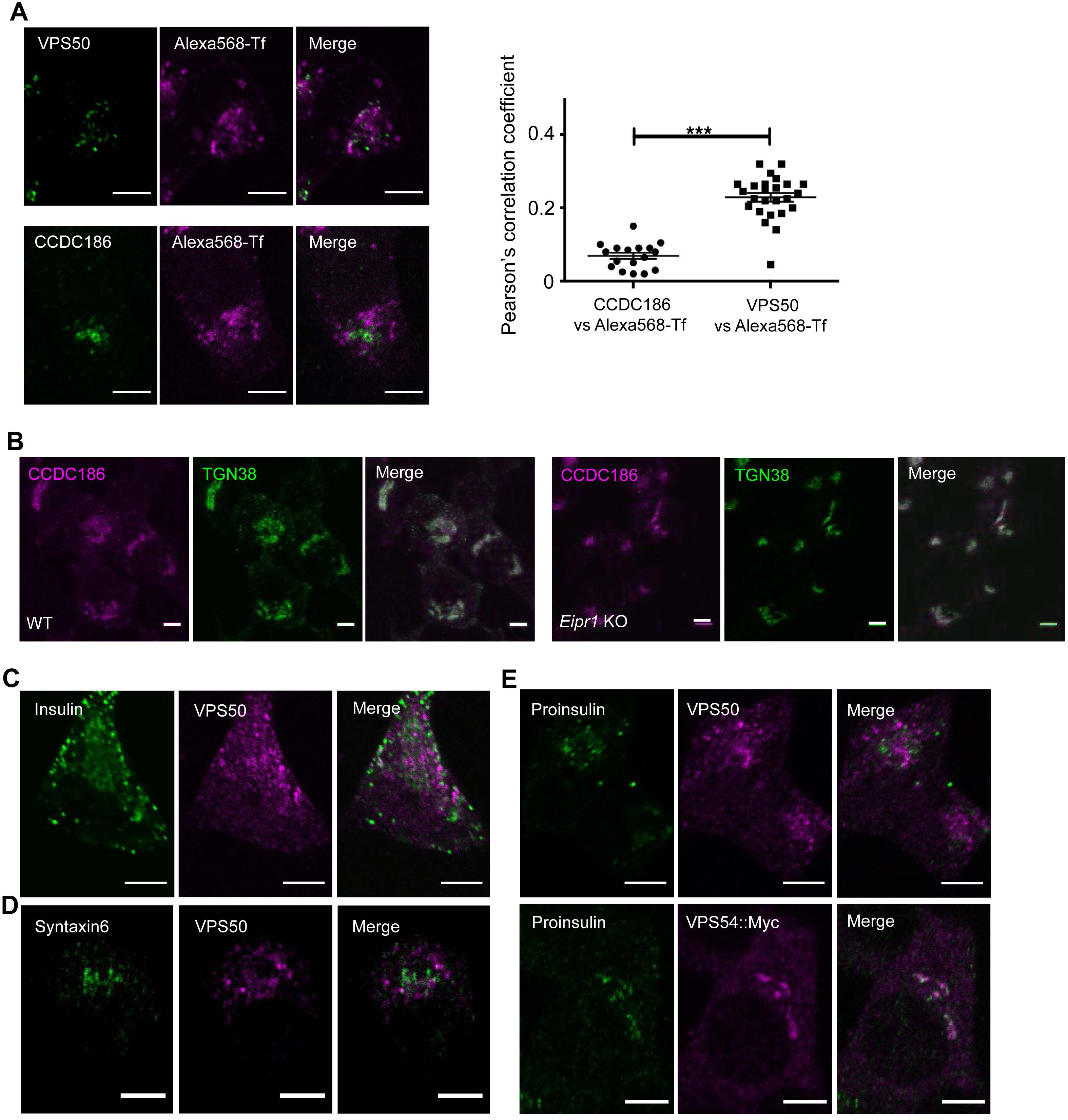
The EARP-specific component VPS50 localizes to two distinct compartments. (A) VPS50 but not CCDC186 partially colocalizes with transferrin. (Left) Representative confocal images of 832/13 cells incubated with Alexa 568-labeled transferrin (Tf) and immunostained for either VPS50 (upper) or CCDC186 (lower). Scale bars: 5 μm. (Right) Pearson’s correlation coefficient was measured to quantify the localization between transferrin and CCDC186 (n=17) or transferrin and VPS50 (n=25). *** p<0.001, error bars = SEM. (B) CCDC186 localization is not altered in *Eipr1* KO cells. Representative confocal images of WT and *Eipr1* KO 832/13 cells immunostained for TGN38 and CCDC186. Scale bars: 5 μm. (C,D) VPS50 only weakly colocalizes with immature and mature DCV markers. Representative confocal images of 832/13 cells immunostained for VPS50 together with syntaxin 6 (iDCVs) or insulin (mature DCVs). Scale bars: 5 μm. (E) VPS50 only weakly colocalizes with proinsulin, but VPS54::Myc largely colocalizes with proinsulin at or near the TGN. Representative confocal images of 832/13 cells or 832/13 cells transiently transfected with VPS54::13Myc and immunostained for VPS50 or Myc and proinsulin. Scale bars: 5 μm.

