## Supplementary figures and images for "EIPR1 controls dense-core vesicle cargo retention and EARP complex localization in insulin-secreting cells"

### Supplementary Figures

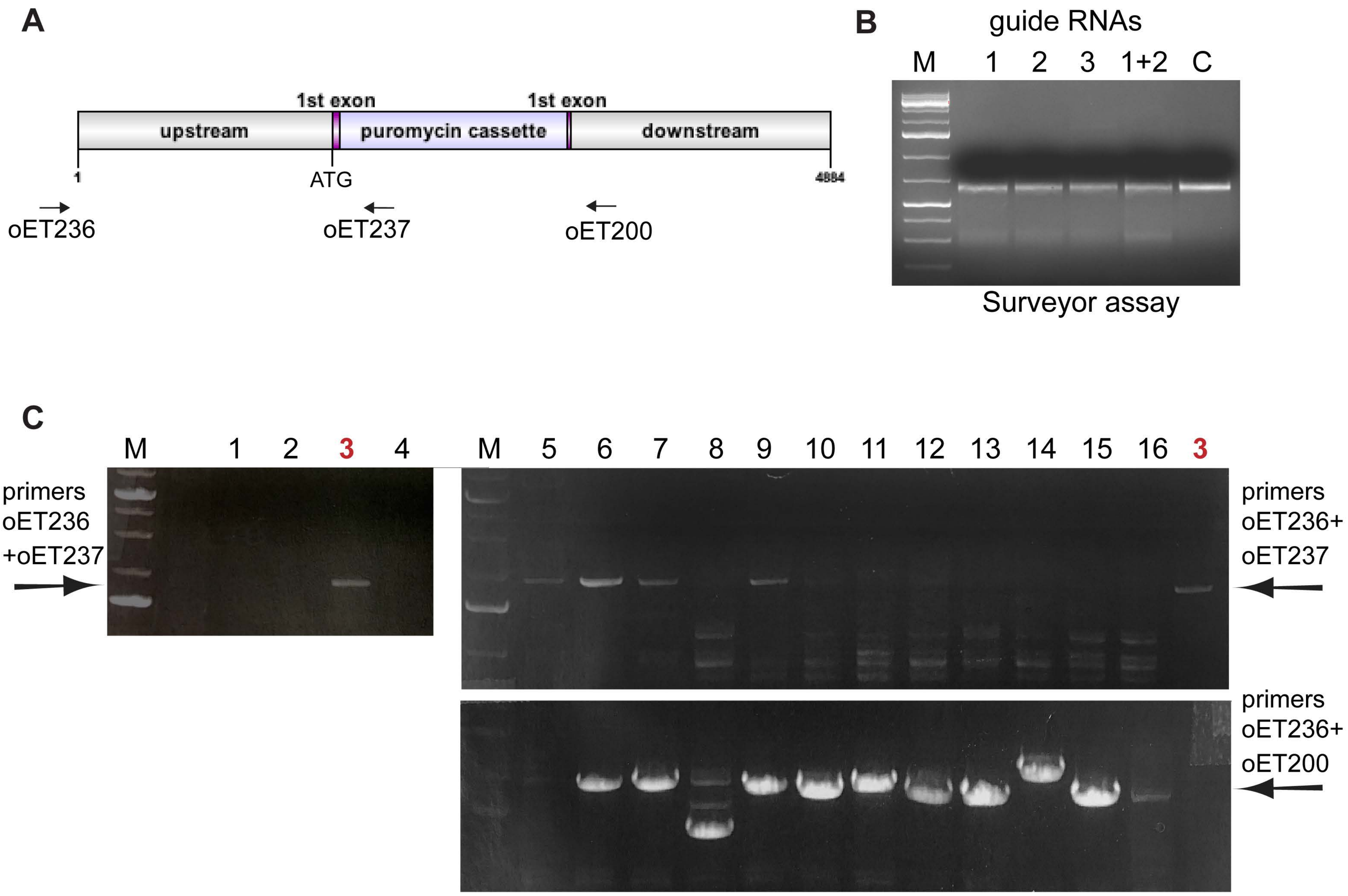

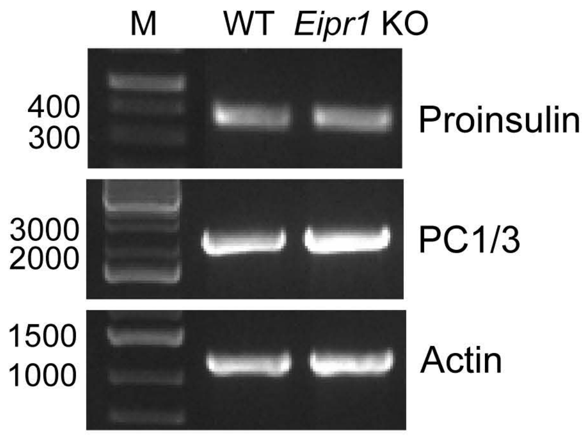

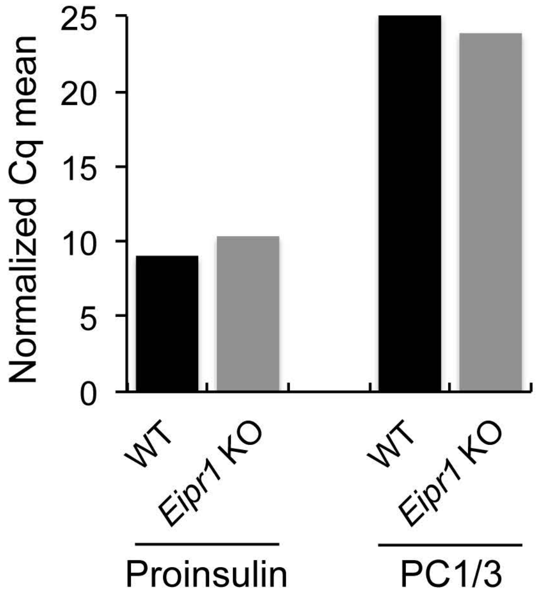

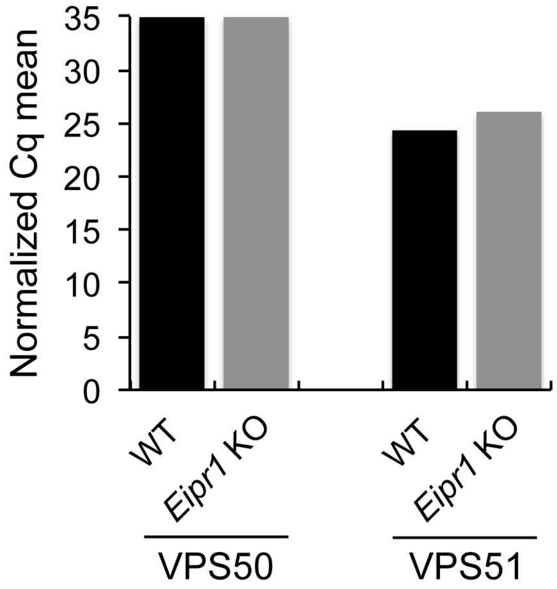

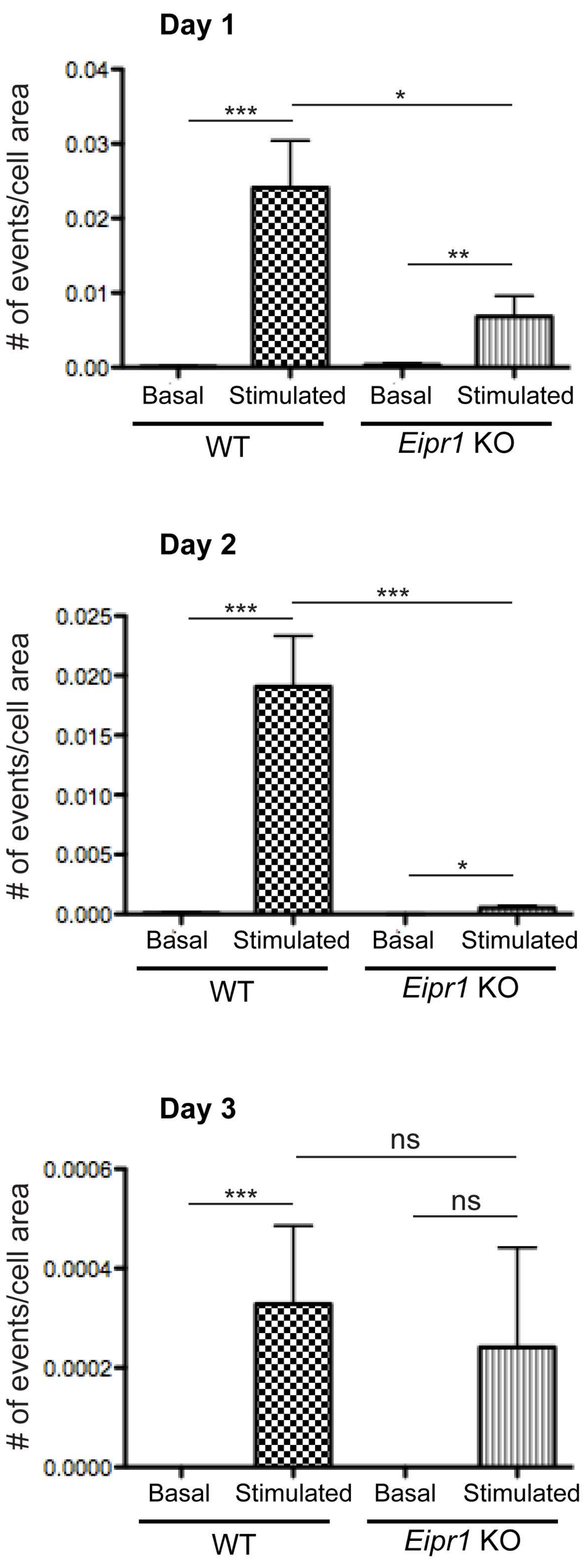

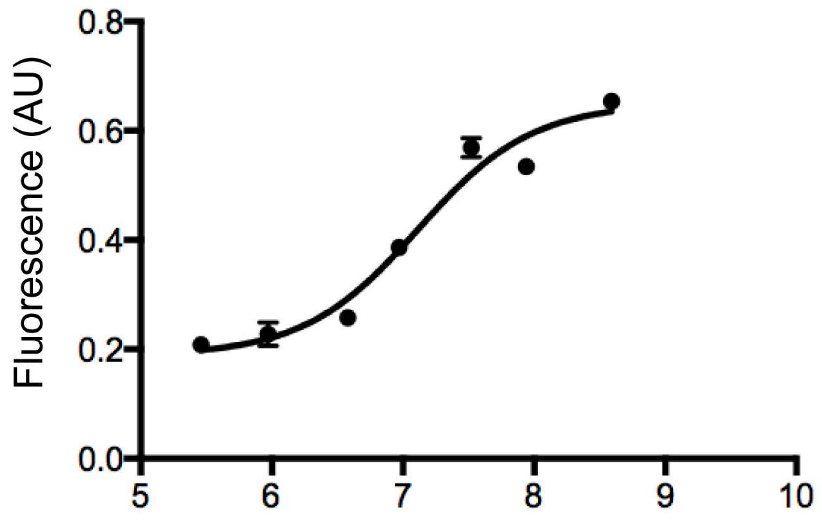

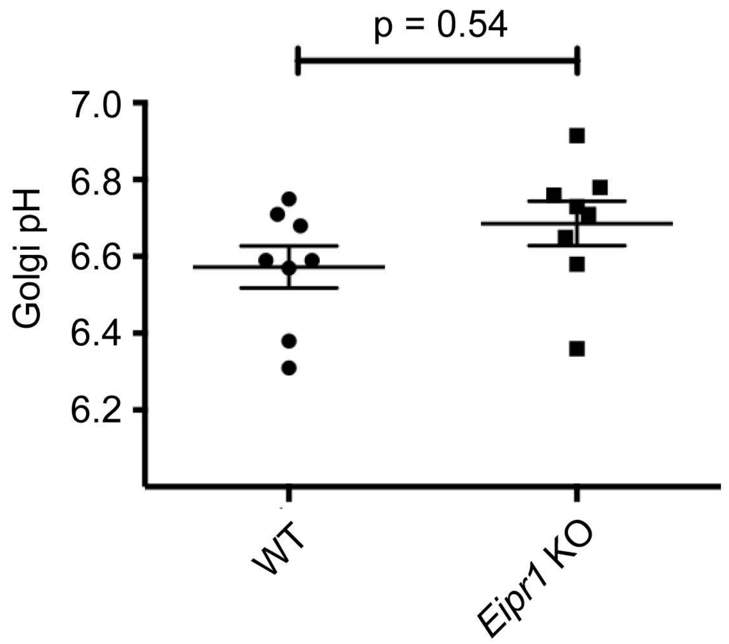

#### Figure S5

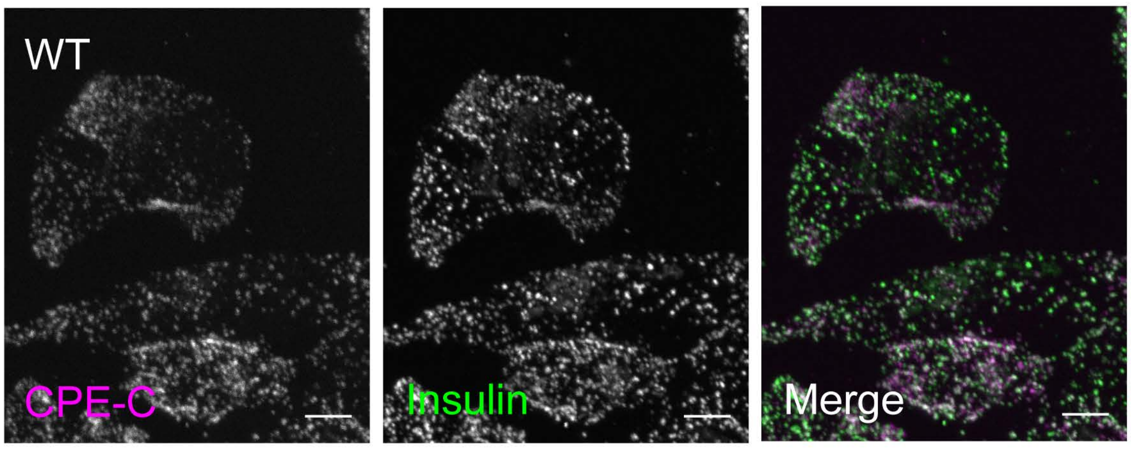

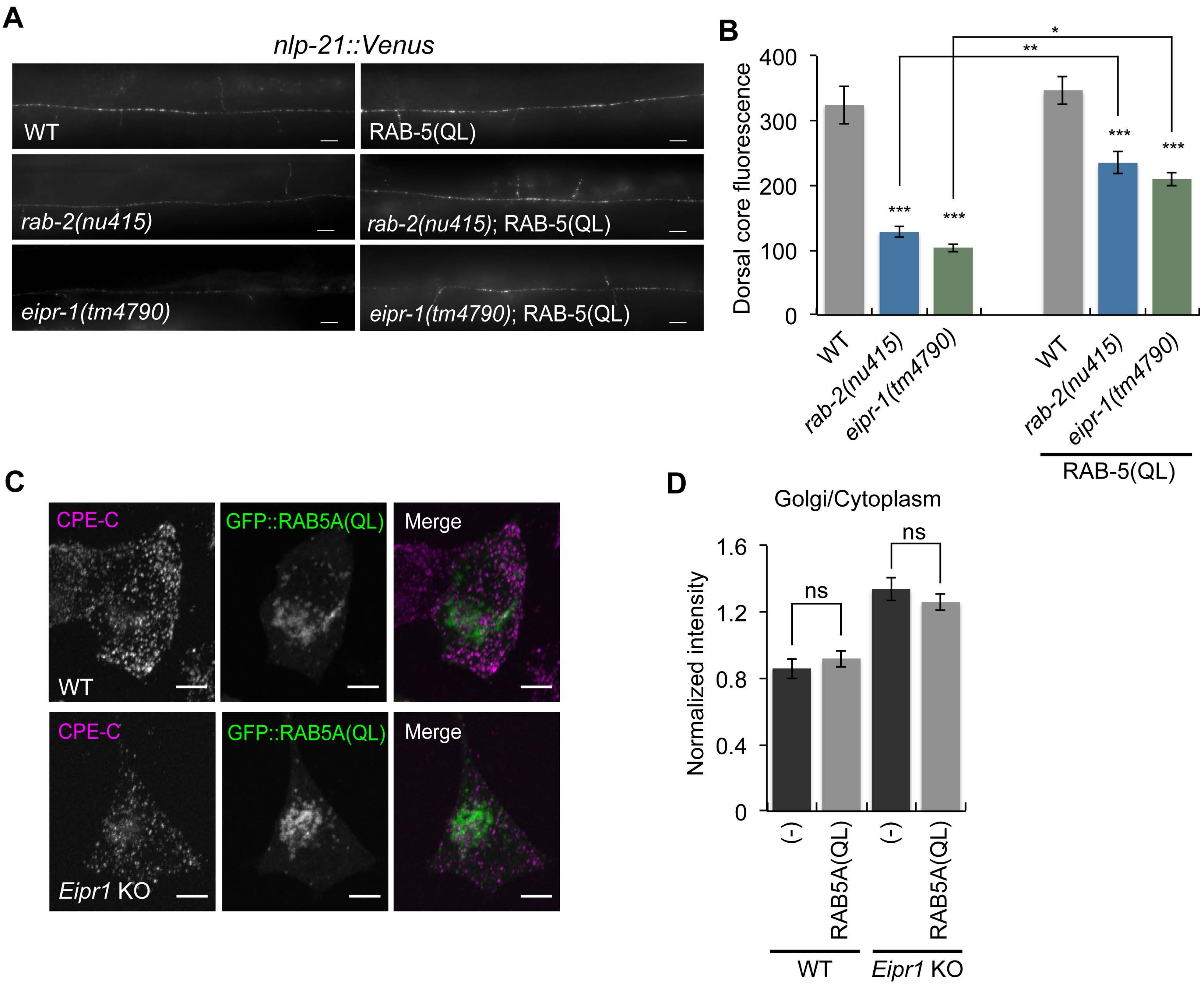

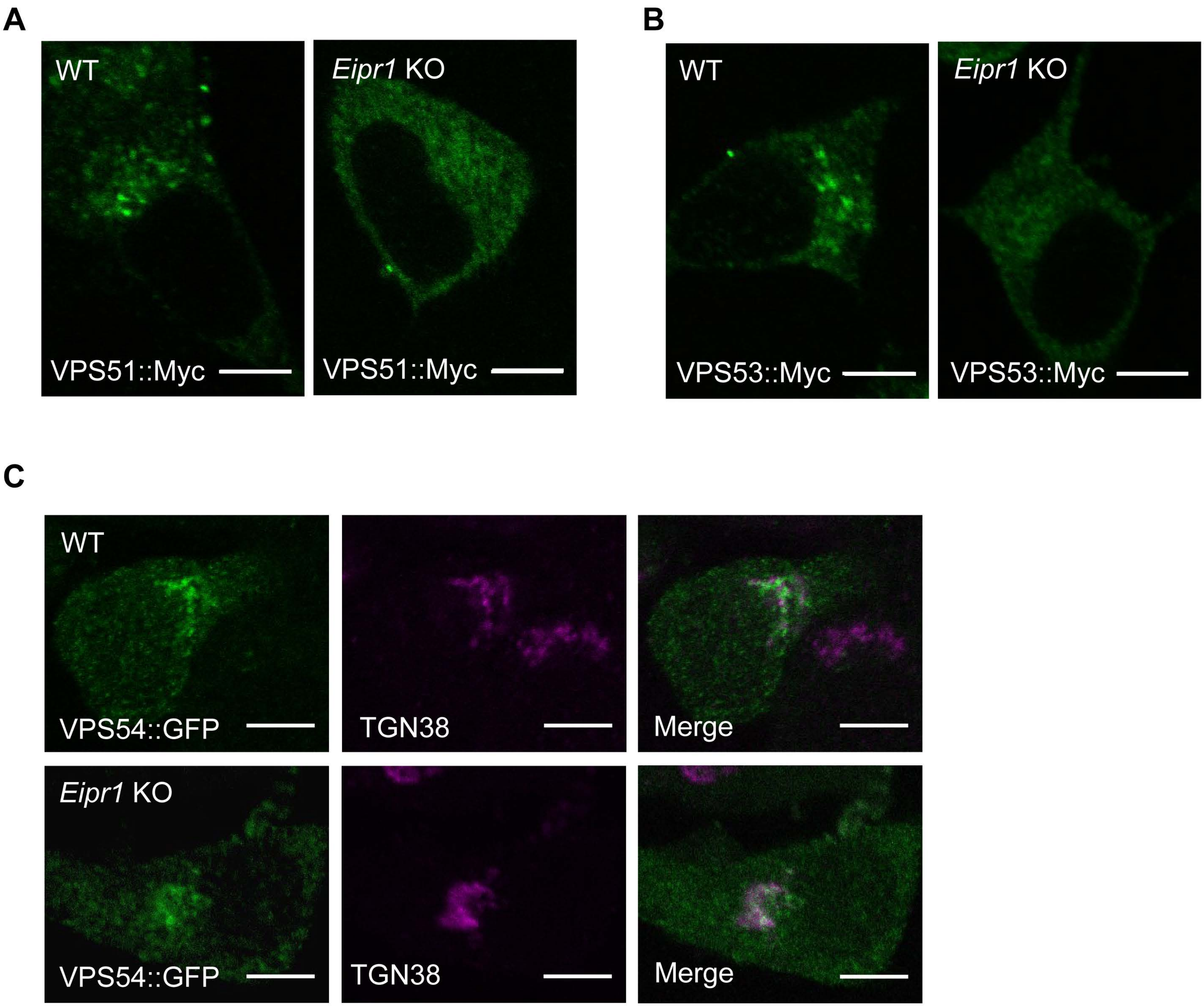

#### Figure S8

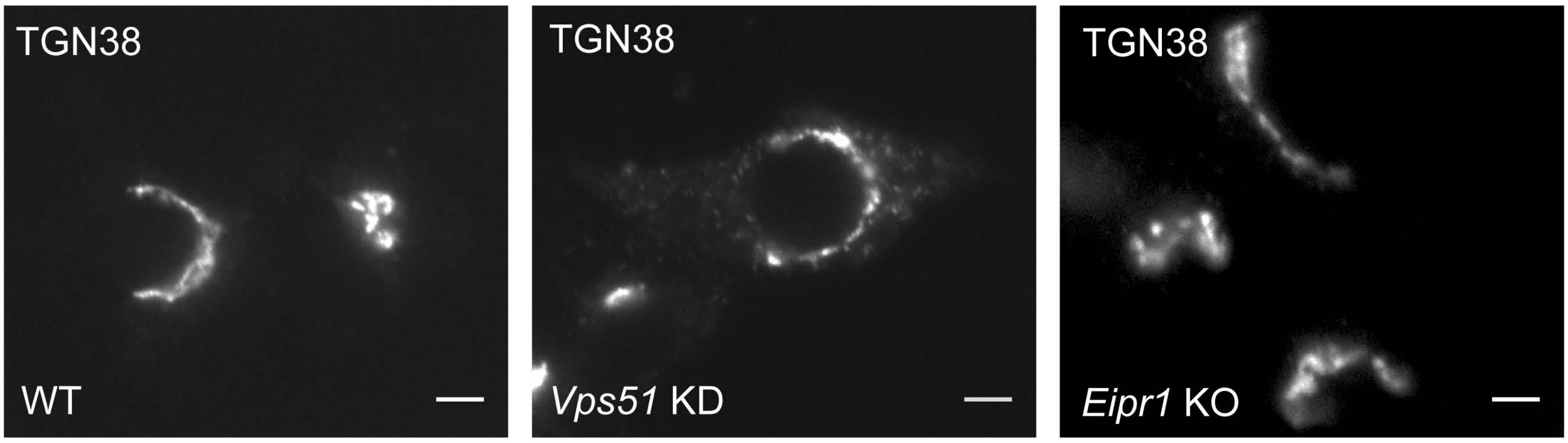

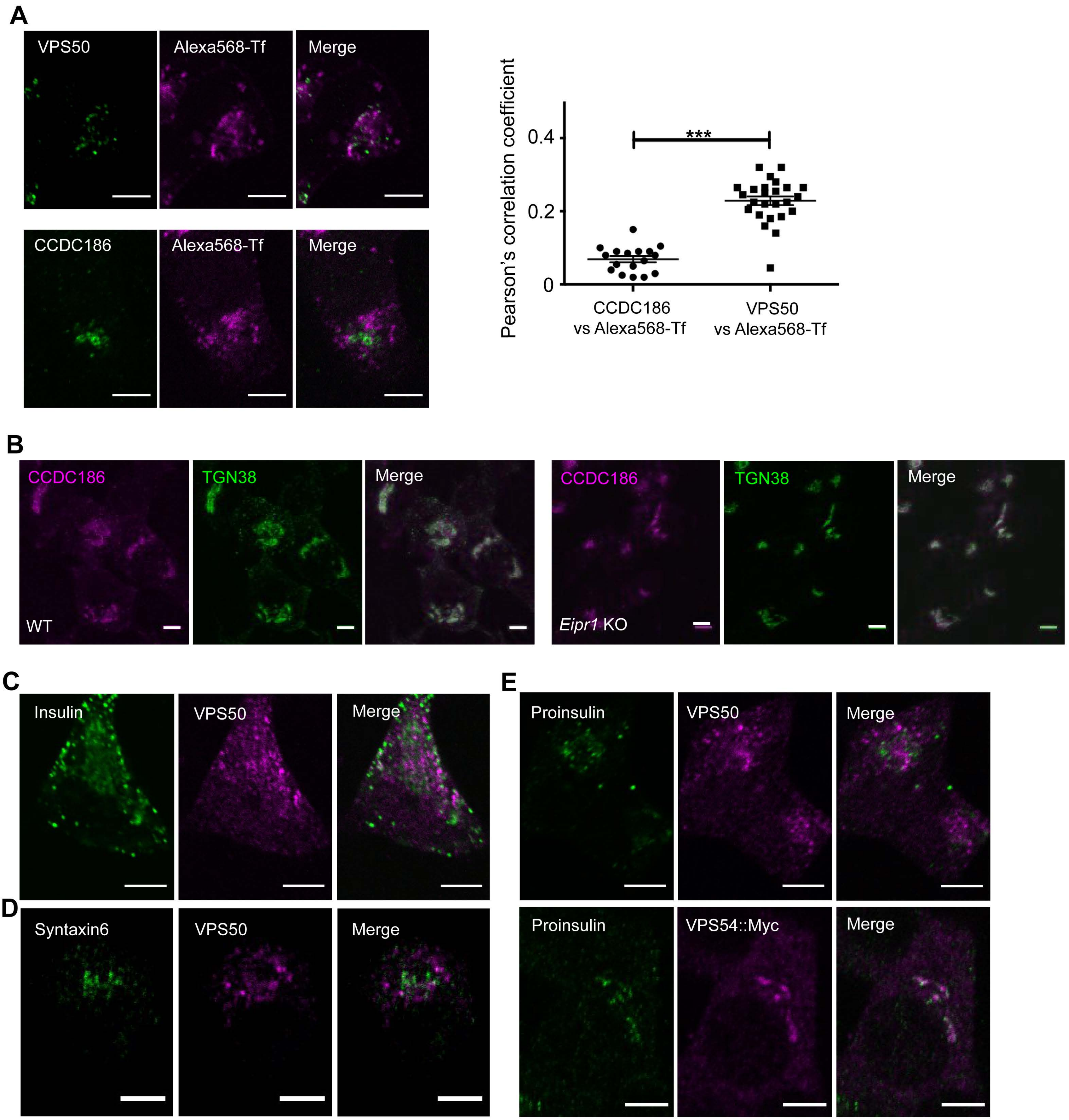
